# Rapid canalization of chromosome conformation-transcription fingerprints during embryogenesis revealed by fully-automated cell identity decoding with CeSCALE

**DOI:** 10.1101/2025.10.22.684035

**Authors:** Konstantinos Ntemos, Fei Xu, Nour-Zaynab Bazzi, Geoffrey Fucile, Hermina Petric Maretic, Ivan Dokmanic, Susan E. Mango, Ahilya N. Sawh

## Abstract

Genome organization into higher-order active and inactive compartments exhibits cell-type specific patterns, which are widely implicated in the regulation of transcriptional activity. During embryogenesis, epigenetic regulation controls cell type specification along cellular lineages with similar transcriptional identities through the coordinated action of chromatin states. However, prevalent single-molecule variability in higher-order chromosome conformation and a lack of precise cell lineage information have previously limited our understanding of the relationship between conformation and transcriptional activity *in vivo*. Specifically, how the conformation-transcription relationship is inherited along cellular lineages through cell divisions is poorly understood. Here, we developed a novel algorithm for cell lineage identification (*C. elegans* Sinkhorn-based Cell ALignmEnt, CeSCALE) combined with single cell genomics to reveal that local conformation-transcription ‘fingerprints’ are associated with, and inherited along the stereotyped cellular lineages of *C. elegans* embryos. Inspired by Optimal Transport theory, CeSCALE provides a fully automated framework for quantifying lineage-resolved individual cell phenotypes *in situ*, across a wide developmental window. Combining CeSCALE with single-molecule chromosome tracing uncovered higher-order interchromosomal block associations, which surprisingly coalesce transcriptionally diverse domains and are independent of lineage identity. Instead, by integrating lineage-resolved chromosome conformations with single-cell transcriptomics, we find that local conformation-transcription spatial relationships (‘fingerprints’), containing both hubs and islands of transcriptional activity, are robustly inherited along lineages. Finally, we find that the canalization of these ‘fingerprints’ represent the rewiring of chromatin states at key developmental stages. Our results suggest that local chromatin environments, but not large-scale compartments, coordinate the dramatically changing transcriptome during embryogenesis.

## 1 Introduction

Animal development is controlled by gene regulation at the epigenetic, transcriptional, and post-transcriptional levels, all intricately orchestrated in space and time. Chromatin factors and epigenetic mechanisms stabilize gene expression programs and canalize cell type identities^1^. One recently appreciated mode of epigenetic regulation arises from the physical arrangement of chromosomal DNA in the nucleus, which can provide a tunable layer of gene regulation during development^2^. Chromatin is organized at multiple interrelated scales: from nucleosomes to loops between regulatory elements, to higher-order structures including domains of contiguous sequences and compartments of non-contiguous sequences^3^. Modulation at each genomic scale provides vast opportunities for gene regulation in response to developmental cues, however compartment-level and inter-compartment level organization is not well understood^4,5^. Originally classified into two broad classes of active (euchromatin) and inactive (heterochromatin) compartments^6–8^, it is becoming clear that the genome folds into complex compartment subtypes of varying genomic size and composition across many species^5^. However, it is poorly understood how higher-order structures are organized *in vivo* at the single-molecule level, especially at the inter-chromosomal level, and if these structures change in a developmental context. Moreover, whether higher-order structures are stably inherited from mother to daughter cells to coordinate cell-type specific gene expression programs in space and time is unknown.

The advent of single cell genomics technologies has improved our understanding of early animal development, tissue composition, and enabled the identification of transient cell states^9–11^. Single-molecule and single-cell analysis of chromosome conformation is critical to improve our understanding since the physical organization of the genome is highly variable regardless of species, scale of chromatin profiled, or method of detection^12–18^. The functional significance of this variability is an active topic of investigation, and one leading hypothesis is that conformational variability is linked to the state or identity of the cell. Supporting this notion, recent work has investigated genome organization at the single cell level and uncovered cell-type specific structural features^15,19–25^, but information on the individual histories of cells in intact animals is lacking. It is currently unclear if structural features can be inherited along cellular lineages to stabilize a particular chromatin state.

Precise knowledge of the cell lineage histories of epigenetic features like chromosome conformation can help explain how these features constrain or facilitate cell functions like transcription at the spatial and temporal levels^26,27^. Do different cell lineages carry different chromosome conformations? How do cell-type specific gene expression programs relate to the prevalent variability in chromosome conformation? These are open questions particularly at the compartment scale, and highlight the need for a synthesized view using single cell multimodal information on chromosome conformation, transcriptional activity, and the collective history and future of all cell divisions. This goal is technically challenging, in particular because of the problem of automated, accurate, and high-throughput lineage identification. However, due to its determinate embryonic cleavage patterns and completely mapped cell lineage, *C. elegans* presents an ideal model to study chromosome conformation-transcription relationships in a developmental context. The current gold standard for cell lineage identification relies on the direct observation and tracing of all cells over time as living embryos develop^28^. Previously developed methods to automate lineage tracing in *C. elegans* embryos rely on tracking the 3D positions of nuclei, where temporal information between consecutive imaging timepoints and observing the cell divisions is required^29–33^. The output of these methods often requires manual curation and editing to produce accurate lineage maps^29,34^. Importantly, it is currently not possible to measure the chromosome conformations and transcriptomes of living cells, therefore a new automated approach is needed to define all cell identities in the embryo from static snapshots of fixed samples.

By exploiting the stereotyped positions of *C. elegans* cells, in this work we developed a novel solution for automated, accurate, and scalable cell lineage identity mapping (CeSCALE) inspired by recent advances in Optimal Transport (OT) theory, which studies how to align two distributions of objects while minimizing displacement costs^35–38^. OT-based methods have recently been successfully used in several other single-cell genomics applications where alignment of cellular features is necessary^39–44^. To the best of our knowledge, CeSCALE is the first method to fully resolve cell lineages in *C. elegans* embryos without the need for tracking cell divisions over time. We combined our lineage mapping method with single-molecule chromosome tracing to discover cell- and lineage-resolved patterns of chromosome compartmentalization through early embryogenesis. Using imaging-based single-molecule and single-cell analysis, we find (1) that chromosomes adopt large interchromosomal block associations, which fluctuate through cell generations and are independent of lineage. (2) We show that these higher-order interchromosomal associations bring together transcriptionally diverse regions of the genome, indicating spatial proximity is not governed by, or a determinant of, shared transcriptional status. (3) On the other hand, we identify local three-dimensional hubs and islands of transcriptional activity, which undergo multiple temporal transitions and are robustly inherited along cellular lineages. (4) Surprisingly, the developmental dynamics of hubs represent large changes in the genomic composition of canonically heterochromatic and euchromatic regions during the zygotic genome activation. Our results suggest the physical arrangement of the genome is in rapid flux to meet the demands of a dramatically changing transcriptome during embryogenesis.

## 2 Results

### 2.1 CeSCALE (*C. elegans* Sinkhorn-based Cell ALignmEnt): mapping fully resolved cell identities through *C. elegans* embryonic development

In *C. elegans*, the relative spatial arrangement of all cells is remarkably consistent across different embryos. This invariant anatomy enables systematic comparison and aggregation of data from equivalent cells (e.g., P4, ABa; see Extended Data Fig. 1, Fig. 1a,b) across many individual embryos to resolve features at single-cell resolution^28^. We leveraged this property to develop CeSCALE (*C. elegans* Sinkhorn-based Cell ALignmEnt), a method to map all embryonic cell IDs simultaneously *in situ*. This approach is grounded in recent advances in Optimal Transport theory, where the key innovation was to formulate the cell identification task as an optimization problem of aligning distance matrices between ground truth reference and test nuclei positions.

**Figure 1:**
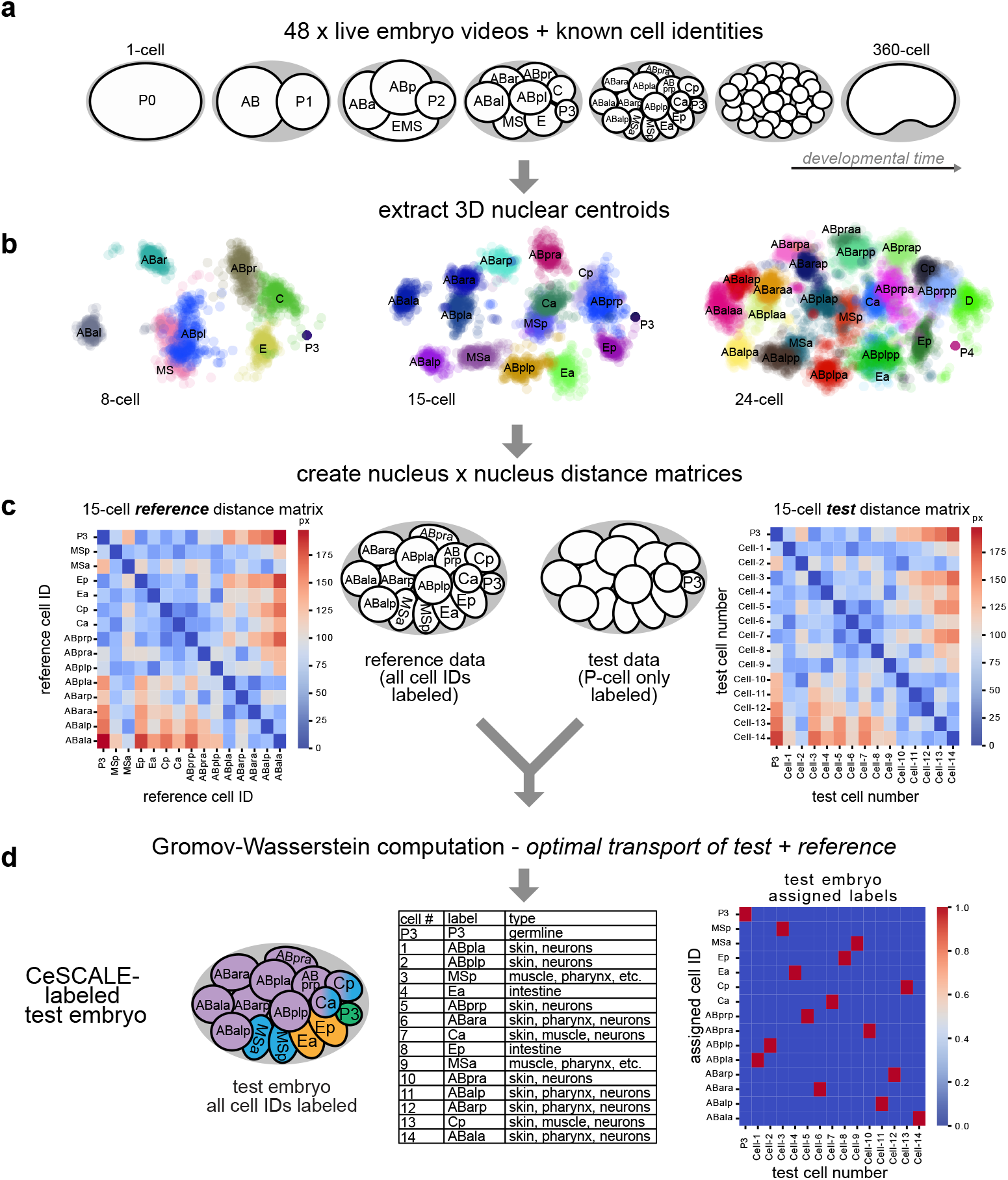
CeSCALE (C. Elegans Sinkhorn-based Cell ALignmEnt) is an automated, scalable framework for cell identification. a) Schematic of *C. elegans* embryogenesis over developmental time. Stages 4-360-cells are covered by our current implementation of CeSCALE. b) Examples of (aligned) point clouds of nuclear centroids from 8-, 15-, and 24-cell embryo live imaging snapshots. Positions are centered on the P-cell. Z-projections are displayed, showing the consistency of spatial positions of the cell identities (e.g., E,C, ABar, etc.) across embryos. c)-d) The pipeline for cell lineage identification using a 15-cell embryo example. Given the all-by-all distance matrices of reference and test embryos, CeSCALE uses optimal transport theory and the known *anchor* cell (P-cell) to predict the unknown cell IDs for the test embryo. By using the obtained transportation plan, we retrieve (by performing maximum weight full matching on the transportation plan) the Cell ID 1-to-1 matching between the cells of the reference map (rows) and the cells of the test embryo (columns). Units of the distance matrices in c) are image pixels, while the assignment matrix (last subfigure) is a permutation matrix and thus, has binary values (0/1).

The first step in CeSCALE involves building comprehensive *reference maps* of cell identities. To create these maps, we used published annotated live imaging data from 48 wild-type *C. elegans* embryos, spanning 2-cell to 360-cell stages, with frames captured every 75 seconds^45–47^ (Fig. 1a). Cell IDs and nuclear centroids (i.e., X, Y, Z coordinates) were extracted using Acetree/StarryNite and manual curation^45–47^ (Fig. 1b). For each embryo image we create the distance matrix containing the distances of all-by-all nuclear centroids. Then, for every developmental age the corresponding reference map is the average (arithmetic mean) of the reference distance matrices, which represent the average spatial configuration of all nuclear centroids. These matrices serve as templates for cell ID labeling of each *test* embryo (Fig. 1c,d), which is unlabeled (i.e., its cell identities are unknown).

CeSCALE performs cell ID assignment by aligning the distance matrix of the given test embryo with the corresponding reference matrix using a Sinkhorn-iteration-based algorithm (Fig. 1c,d). Through this process, we can rearrange the rows and columns of the one matrix to make it as similar as possible to the other (by minimizing an appropriate cost function). To perform the alignment, we extended the algorithm originally proposed to compute the Gromov-Wasserstein (GW) distance between distance/similarity matrices in Optimal Transport problems^36^, in order to capture the nuances in our setup. More specifically, to maximize the quality of cell alignment and reduce ambiguity in cell ID predictions, we extended the GW computation to include a known position of a single anchor cell in each test embryo. We chose the germline precursor cells (P-lineage: P0, P1, P2, P3, P4, Z2/Z3, see Extended Data Fig. 1) since they are present throughout embryogenesis and can be easily detected experimentally. Our reformulated optimization problem fixes the P-cell position and uses projected gradient descent with Sinkhorn projections (as in ^36^) to obtain the transportation plan which obtains the cell-to-cell mapping, and thereby predict all the remaining cell IDs for test embryos (Fig. 1d). Detailed algorithm descriptions, mathematical formulations and derivations are described in the Methods section. We note that our methodology can be extended with appropriate modifications to using an arbitrary number of cells as anchors.

### 2.2 CeSCALE achieves >94% average accuracy in cell identification over a broad developmental window

We evaluated CeSCALE’s accuracy by comparing predicted cell identities in test embryos to the ground truth labels. Reference data contained all labels to make reference maps, while test data included only the P-cell label and they were used to compute the accuracy of the predicted labels. The average accuracy for every embryo image was computed in a 1-vs-all fashion, where the embryo under investigation was the test embryo and the reference map was constructed from the rest of the embryos (see Methods). The accuracy for each image was calculated as (1-(N_errors_/N_total_))*100, where N_errors_ is the number of incorrectly labeled cells (i.e., wrong cell ID assigned to them) and N_total_ corresponds to the total number of cells. A value of 100% accuracy means that correct cell IDs were assigned to all cells in the image. CeSCALE achieved >=94.7% accuracy (averaged across all available embryos of a particular stage) for all stages up to the 360-cell embryo (Extended Data Fig. 2a), except for the 4-cell embryo which is 84% (see below).

At the individual cell level, we observed that the average cell accuracy (average across all available cells in all ages and all embryo images) is remarkably high (97.09%), including near-perfect performance for most individual cell IDs (Fig. 2a). Uniquely for the 4-cell embryo, ABp and EMS cell accuracy is lower than other cells through development (68%). This is due to the symmetry of the 4-cell embryo, where distances between ABp and EMS to all other cells are similar. For later stages, this embryo symmetry breaks and the accuracy is significantly higher across all available cells (Fig. 2a, Supplementary Data 3). We note here that CeSCALE is not restricted to a specific embryonic stage, but the available annotated data used to verify the algorithm accuracy spanned the 4 to 360-cell stages (see Methods and Supplementary Data 1).

**Figure 2:**
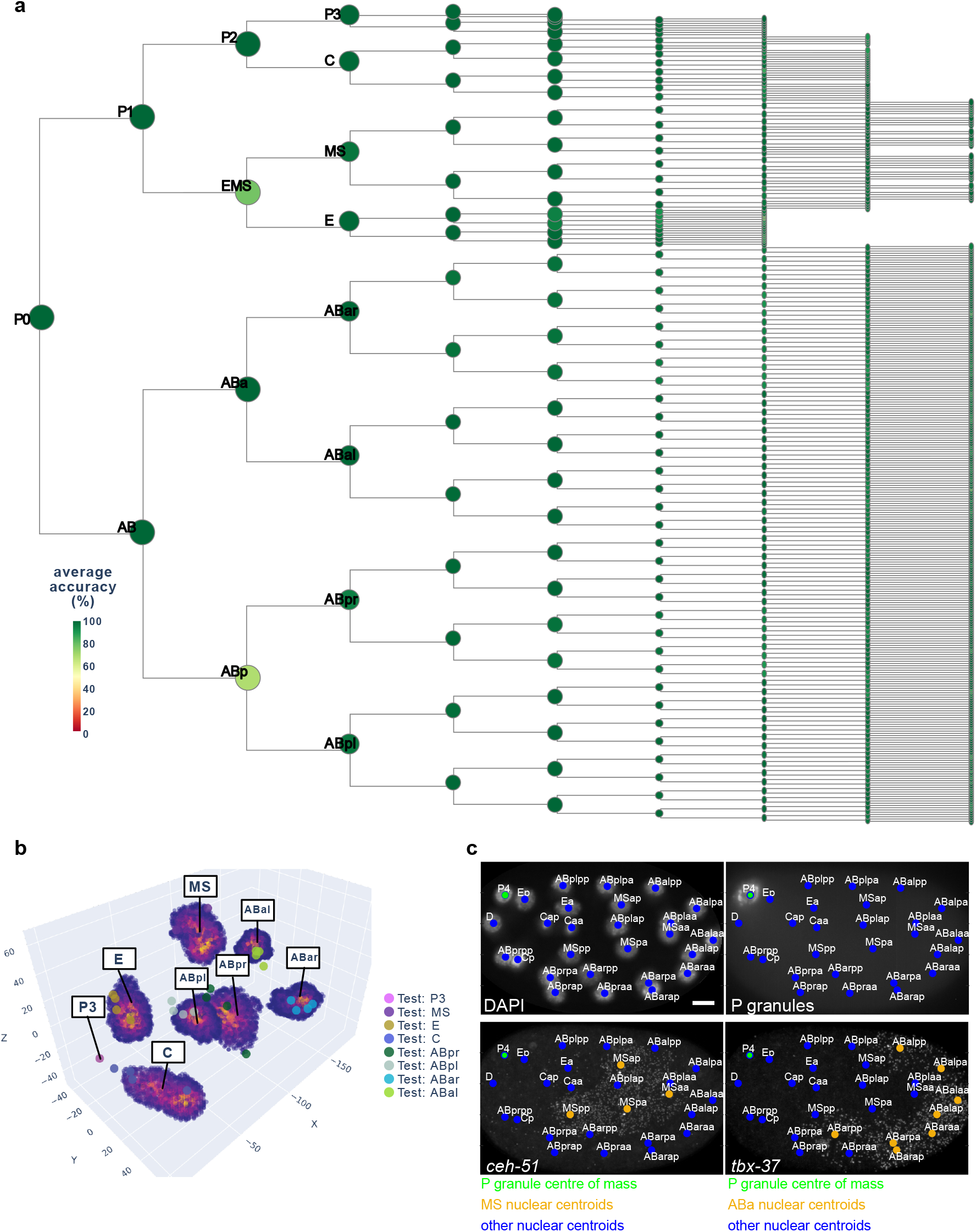
CeSCALE achieves high accuracy in cell identification from live and fixed images over a broad developmental window. a) Dendrogram of lineage relationships between cells from the 1-cell to 360-cell stage annotated with the CeSCALE accuracy from live images. 1-cell and 2-cell embryos do not require CeSCALE for label prediction since the P-cell marker identifies them with 100% accuracy (Extended Data Fig. 1). b) KDE point clouds of nuclear centroids from reference 8-cell embryos, overlaid with test embryo positions from fixed images (4 embryos), showing that cell IDs from fixed embryos fall within the corresponding expected reference areas, thus indicating both accurate cell labeling via CeSCALE and consistency in the spatial cell arrangements between the test embryos and the reference embryos. Positions are centered on the P-cell. See Supplementary Data 6, 7, 8 for additional stages in interactive plots ( 8-, 15-, 24-cell stages respectively). c) RNA smFISH overlay of CeSCALE predicted cell IDs in fixed embryos. *ceh-51* foci mark the cytoplasm and nuclei of MS-cells, while *tbx-37* foci mark the cytoplasm and nuclei of ABa cells. Nuclear centroids are labeled with coloured circles. Maximum intensity z-projections are displayed. Scale bar, 5 µm.

The measurement of many cellular features, including chromosome conformation, cannot be performed in living samples since they preclude the penetration of reagents like antibodies and oligonucleotide probes. Thus, samples need to be fixed and permeabilized, which could lead to changes in the embryo shape and cellular positions. To test if CeSCALE can reliably label cell IDs in images from unseen fixed samples, we applied the algorithm to fixed embryos where we defined the P-cells with a fluorescent protein marker localizing to perinuclear germ granules (PGL-1::GFP at the endogenous locus). Embryos ranging from the 4-cell to 93-cell stage were immunostained with a GFP antibody and imaged with widefield fluorescence. Nuclear centroids were obtained from the 3D image stacks using a watershed-based segmentation pipeline, P-cell nuclei were automatically identified using the GFP signal (see Methods, Extended Data Fig. 3), and CeSCALE was used to process these coordinates and label all somatic cell IDs. As ground truth labels are not available for fixed samples, the quality of labeling was evaluated in three alternative ways: 1) Procrustes alignment per embryo, 2) confidence score of each nucleus, and 3) RNA single molecule FISH of marker gene transcripts.

First, to evaluate the overall spatial displacement of all predicted labels per embryo with respect to the reference configuration, we first obtain the cell labels with CeSCALE and then we super-impose the test embryo cell positions over the reference cell configuration via the Procrustes algorithm^48,49^ (in order to find the optimal rotation, translation and scaling for the test embryo) and calculate the normalized Procrustes distance. This metric is defined as the normalized sum of squared residuals between the super-imposed nuclei positions of reference and test embryos, (see Methods) and expresses a goodness-of-fit measure of the overall alignment (the smaller the distance the better alignment). We observe that the normalized Procrustes distance is close to 0 for all embryos, implying good alignment with small overall displacement, for both live (average value 0.02) and fixed embryos (average value 0.06) (Extended Data Fig. 2b).

The normalized Procrustes distance is designed to capture the global alignment of the embryo while ignoring possible local deviations of specific cells. As a complementary, cell-level metric, we compute a per-cell-identity confidence score: the normalized likelihood of the observed nuclear centroid under the spatial distribution of the corresponding reference cell ID. The spatial probability distribution for every cell ID is approximated by performing Kernel Density Estimation (KDE) on the nuclear centroid point clouds of the reference cohort (see Methods) (Fig. 2b). This per-cell ID confidence score takes values between 0 and 1, with 0 implying that the cell position lies outside the distribution of the corresponding cell ID expected positions, while a value closer to 1 indicates agreement with the expected spatial configuration. In fixed embryos, per-cell confidence is high (average per-cell confidence across all available cells and images is 0.84; Extended Data Fig. 2c), thus implying robust alignment. We note that this metric is very conservative, where a low score could be due to a nucleus lying just outside the KDE point cloud (e.g., in the case of recent cell division). Third, we performed single-molecule RNA FISH on embryos for lineage-enriched transcripts. We probed for *ceh-51* and *tbx-37*, which are known to be enriched in the MS and ABa cells respectively^50–52^. Comparison of CeSCALE-labeled embryos with the distribution of RNA foci confirmed that predicted MS and ABa nuclear centroids were embedded in cells expressing *ceh-51* and *tbx-37* respectively (Fig. 2c), indicating accurate cell ID prediction. Together, these three measurements—Procrustes distance, per-cell-ID confidence, and RNA FISH—demonstrate that CeSCALE accurately predicts cell identities in fixed embryo images.

All together, these results demonstrate that CeSCALE can be applied to virtually any live or fixed image frame from embryos in a broad developmental window to automatically and accurately identify all cell IDs in a robust way. The proposed method for cell labelling is simple and intuitive (based only on the distances between nuclei) and highly scalable due to the use of the Sinkhorn algorithm^35,53^.

### 2.3 Combined CeSCALE and single-molecule chromosome tracing identifies global lineage differences in chromosome conformation

To directly measure higher-order chromosome conformation at the single molecule level *in situ* during development, we employed chromosome tracing by multiplexed sequential FISH^12,13,54^ on two entire autosomes simultaneously, totalling ⅓ of the *C. elegans* genome (chrI and chrII, see Methods). Chromosome tracing probes targeted the central 100kb genomic regions of 14 domains on each autosome (Fig. 3a-c, Extended Data Fig. 4) and uses a dual-probe design, where primary FISH probes coat entire chromosome territories, and tiled region-specific secondary probes detect numerous individual genomic regions along chromosome lengths in iterative sequential hybridizations on the same sample. We collected data from embryos ranging between the 4-cell and 93-cell stage, to a total of 4775 single-molecule chromosome traces. This age range captures the birth and specification of all cell lineages, the establishment of the embryo’s anterior-posterior asymmetric and left-right symmetric axes, gastrulation, and the minor and major bursts of the zygotic genome activations^55–57^ (Extended Data Fig. 1). On average, chrI and chrII displayed relatively smaller intra-chromosomal distances between domains than inter-chromosomal distances (Fig. 3d), consistent with previous bulk Hi-C data on mixed-stage embryos^58–62^. However, all inter-domain distances displayed high standard deviation, with only slightly lower standard deviation between contiguous domains (Fig. 3d, right). These data indicate that individual chromosomes differ markedly in their conformations both within and between chromosomes, consistent with previous single-molecule results on intrachromosomal compartments in pre-gastrula embryos^12,13^.

**Figure 3:**
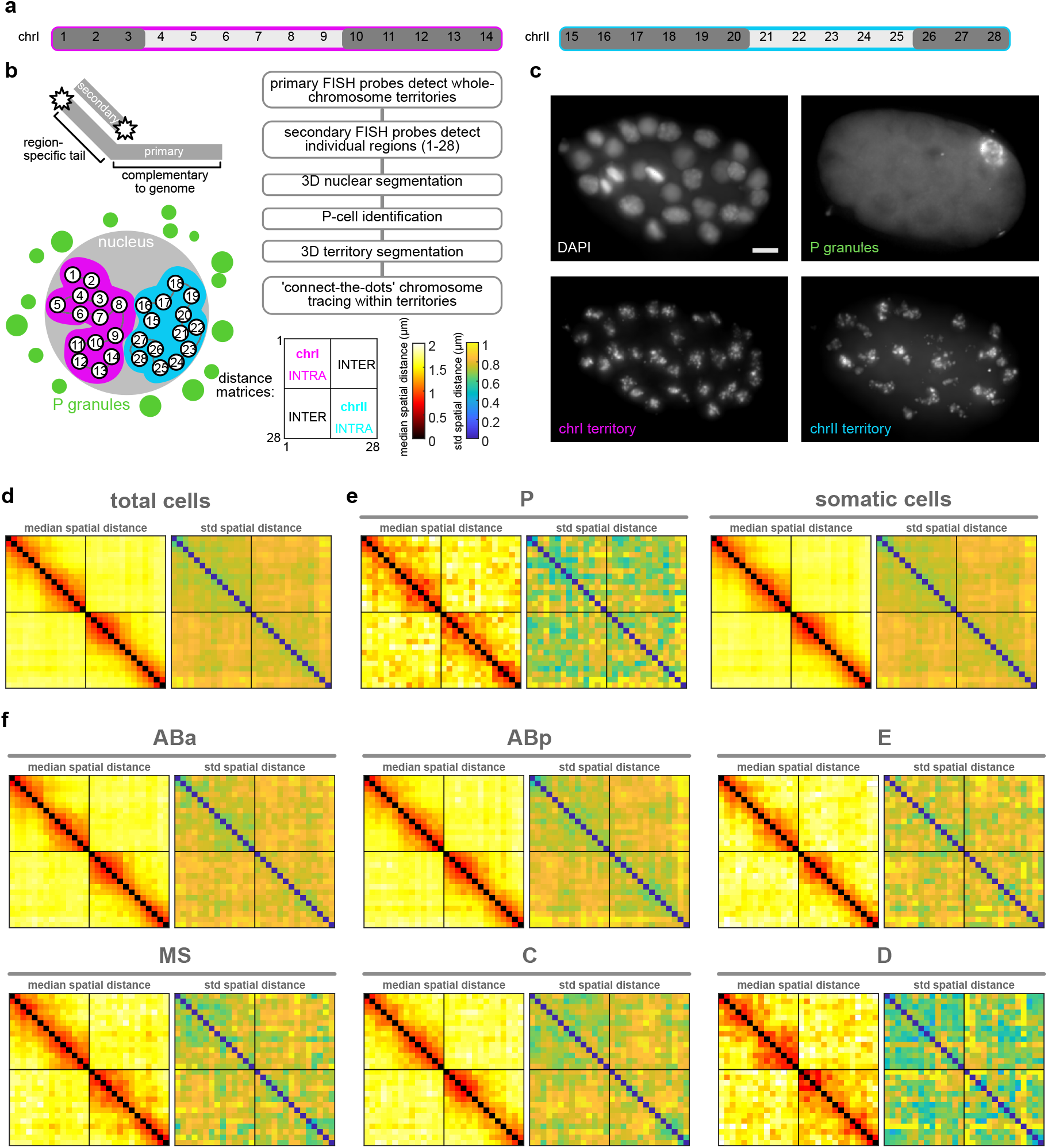
Combined CeSCALE and chromosome tracing identifies P and D lineage cells have more compact chromosome conformation than anterior cells. a) Schematic of regions probed during chromosome tracing experiments. Regions 1-14 span chrI and regions 15-28 span chrII. Dark gray regions correspond to chromosome arms, light regions correspond to chromosome centres. b) Chromosome tracing experimental and data processing overview. c) Example images of chromosome territories, nuclear volumes, and P-cell. Maximum intensity z-projections are displayed. Scale bar, 5 µm. d) Median spatial distance matrix (left) and standard deviation of spatial distance matrix (right) for all chromosome traces detected in embryos. n= 4775 traces. e) Distance matrices of traces in P-cells (n= 193 traces) vs. somatic cells (n= 4582 traces). f) Distance matrices of traces in ABa (n= 1540 traces), ABp (n= 1557 traces), E (n= 441 traces), MS (n= 481 traces), C (n= 448 traces), and D cells (n= 115 traces). Colormap scales as in b) for all matrices.

We hypothesized that the observed conformational variability could be due to both inter- and intra-lineage heterogeneity. Chromosome traces originating from the P-lineage displayed more compact conformations with increased inter-chromosomal proximity (Fig. 3e) and lower standard deviation compared to the total or all somatic traces (Fig. 3d,e). High compaction and low variability in the P-lineage could be linked to the overall transcriptional silence of these cells, compared to the somatic lineages which are transcriptionally active^55,57^. To identify the patterns found among the somatic lineages, we identified all cell IDs with CeSCALE (Extended Data Fig. 5), and automatically assigned somatic traces to cell IDs based on their bounding nuclei (see Methods). When traces were parsed by lineage, the standard deviation of spatial distances decreased compared to pooled somatic cell traces (Fig. 3e, f). Similar to the P-lineage traces, D-lineage traces displayed high compaction at the intra- and inter-chromosomal level, as well as low standard deviation (Fig. 3f, Extended Data Fig. 6). This similarity could be due to the fact that the D-lineage is closest in lineage history to the P-lineage (Extended Data Fig. 1), indicating that these posterior cells may maintain a memory of overall chromosome conformation across cell divisions. The other somatic lineages displayed more subtle differences in intrachromosomal conformation and higher standard deviations than P and D-lineages (Fig. 3e,f, Extended Data Fig. 6). Highest variability was observed in the AB-lineage, which is also the largest population of cells and traces (Fig. 3f), suggesting that intra-lineage heterogeneity also strongly influences chromosome conformation.

### 2.4 Whole chromosomes adopt higher-order conformations with prevalent territory intermingling, in proportions independent of lineage distance

An outstanding question from previous analyses of single-molecule chromosome conformations^12,13,63^ is whether different classes are associated with different cell types or lineages. To test this, we used our previous unsupervised clustering method to unbiasedly identify prominent sub-populations among our single-molecule data^12,13,63^ (see Methods). Using all pairwise distances between all 28 regions of chrI and chrII as the features to cluster individual traces (378 variables), we identified 11 broad classes of conformations (see Methods, Fig. 4a). Each of the 11 clusters significantly differed at unique regions in their pairwise distance patterns, compared to the remaining pool or other clusters (Extended Data Fig. 7a, 8), demonstrating they are *bona fide* sub-populations with different folding patterns.

**Figure 4:**
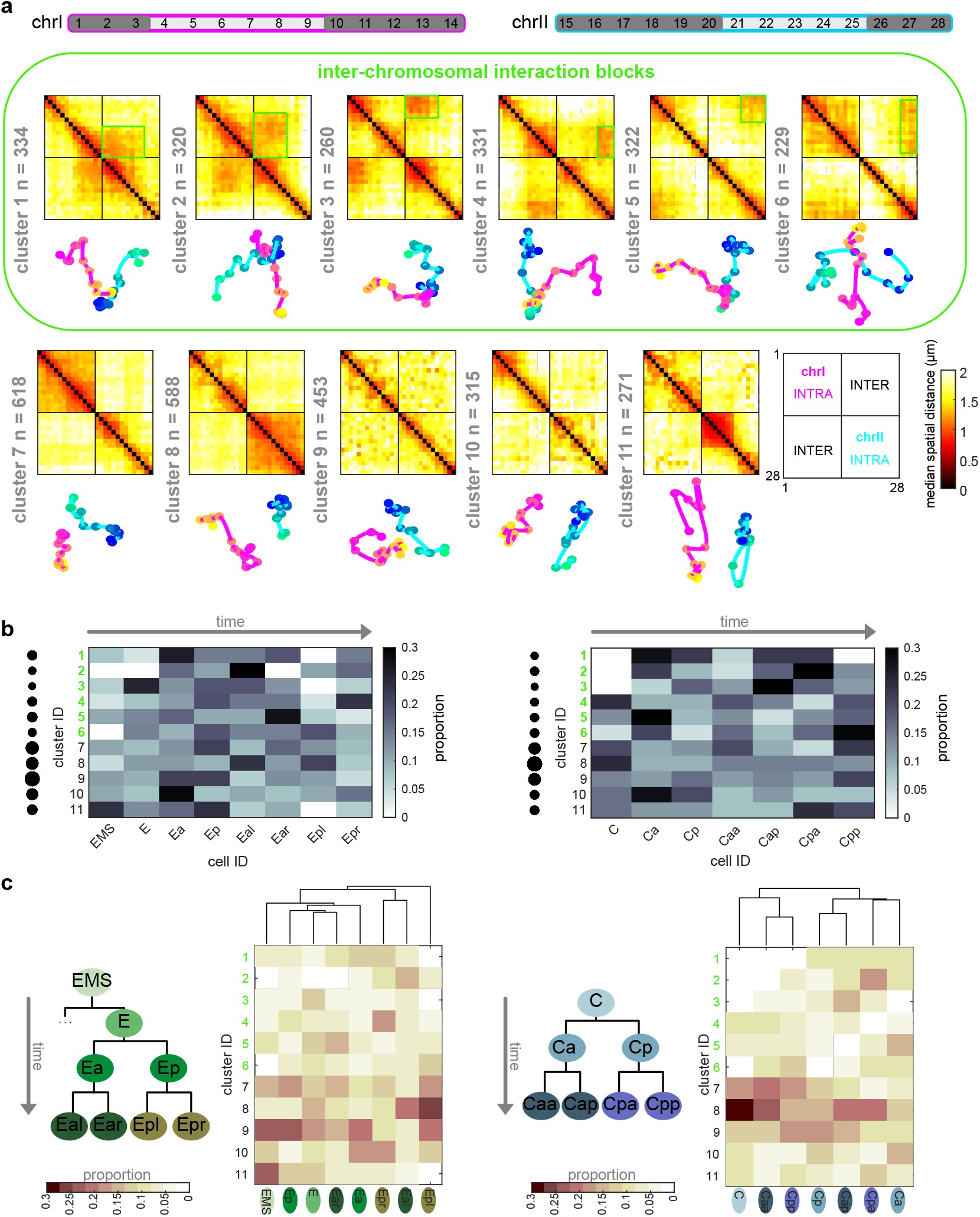
Higher-order interchromosomal blocks characterize early embryonic chromosome conformation lineage-wide. a) Median distance matrices and 3D models of chromosome clusters. Clusters 1-6 display prominent territory intermingling in interchromosomal interaction blocks (green outlines). b) E- and C-lineage examples of cluster proportions. To determine in which cells each cluster is enriched, each matrix row is normalized to the total amount of traces per cluster. c) E- and C-lineage examples of cell proportions. To determine the proportion of clusters within a cell, each matrix column is normalized to the total number of traces per cell ID. The columns of the matrix are ordered using hierarchical clustering by Euclidean distance.

Previous population-based analyses have shown that chromosome territories are largely distinct, with inter-chromosomal regions possessing low contact frequency^58–60,62^. Compared to intra-chromosomal contacts, relatively rare inter-chromosomal contacts are proposed to occur primarily along the edges of territories^64,65^. Surprisingly, our analysis of individual chromosome conformation variability identified large blocks of high inter-chromosomal proximity (Fig. 4a), indicating that chrI and chrII territories display prevalent intermingling *in vivo*. None of the clusters closely resembled the average conformations of each lineage (Fig. 3e,f), demonstrating the different classes were obscured by averaging. Inter-chromosomal interactions were not limited to edge-edge territory proximity, as in cluster 9 and previous studies, but were often found encompassing up to about half of the chromosomes (clusters 1-6, Fig. 4a). These inter-chromosomal interaction blocks contained lower standard deviation values compared to their surroundings, in some instances lower than intra-chromosomal pairs within a structure (Extended Data Fig. 7b). The high degree of single-molecule conformational variability observed explains why these blocks were previously missed when considering average whole-embryo or whole-lineage data from both imaging (Fig. 3) and sequencing-based methods^58–62^. Overall, clusters 1-6 show that non-homologous chromosomes often fold together in particular genomic blocks during embryogenesis (Fig. 4a), in associations as close or closer than intra-chromosomal regions, opening up the possibility for regulation at this scale.

If higher-order conformation is inherited in cell types from mother to daughter cells, we expect that cells with close lineage history will have similar conformations. We therefore asked how the clusters varied in cells as development progresses in different lineages. Considering all stages aggregated, the different lineages displayed subtle differences in the prevalence of clusters between and within lineages (Extended Data Fig. 9a). However, when proportions were binned by cell ID to track patterns over developmental time, different patterns emerged among cells in all cases (Fig. 4b-c, Extended Data Fig. 9b). Examining each cluster’s cellular origins, we found that each cluster appeared in each lineage, albeit in different proportions in different cells (Fig. 4b, Extended Data Fig. 9b). For example, cluster 5 was found in its highest proportions in Ear, Ca, and D compared to cells of the same lineage, but was also detected in every other cell (5% or greater). The prevalence and timing of interchromosomal interactions (clusters 1-6) also fluctuated among cells of the same lineage (Fig. 4b, Extended Data Fig. 9b). For example, clusters 1-3 appear in high proportions in the C lineage only after the birth of Ca/Cp (15-cell), but are present in low proportions in the other lineages from the earliest stages probed (4-cell)(Fig. 4b, Extended Data Fig. 9b). All higher-order conformations are therefore produced in all cell types throughout the embryo’s life.

Examining the conformational makeup of individual cells, we found that sister cells, which are likely to exhibit the greatest molecular similarity, did not carry the closest conformation patterns (Fig. 4c, Extended Data Fig. 9b). For example, sisters Epr/Epl and Cpp/Cpa cells differed markedly in cluster proportions, as shown by hierarchical clustering of the conformation patterns (Fig. 4c). Broadly, this observation held true for all lineages, showing that lineage distance does not dictate large-scale chromosome conformational differences in cells of the early embryo (Fig. 4c, Extended Data Fig. 9b). These results suggest either conformations emerge *de novo* at each cell division, similar to previous observations of chromatin-lamina association^66^, or that sister cells can have an asymmetric memory of conformation from their mother cell. Moreover, since different lineages carry similar conformations (i.e. all clusters appear in significant proportions across all lineages), these data suggest similar higher-order conformations are conducive to different cellular functions.

### 2.5 Higher-order interchromosomal blocks coalesce transcriptionally diverse regions

The higher-order chromosome conformations identified through our clustering approach did not correlate with cell lineages in the embryo, suggesting that they are not coupled to lineage-specific transcriptional programs. Higher-order chromosome compartments are thought to spatially coordinate active or inactive chromatin to co-ordinate gene expression^4,5^, but these structures are not well studied at the single molecule level *in vivo*. We therefore asked whether conformations coordinate uniform transcriptional activity or, as indicated by our lineage analysis, similar conformations are conducive to different gene expression outputs and cellular functions. Several recent studies have identified the lineage-resolved transcriptomes of embryonic cells using single-cell (sc)RNA-seq^50–52,67^. To determine how local transcriptional activity of different genes during the zygotic genome activation (ZGA) intersects in 3D, we integrated scRNA-seq data with our lineage-resolved chromosome tracing.

We first compared the transcriptional landscapes of chrI and chrII conformations (clusters 1-11), across the major cellular lineages of the embryo (ABa, ABb, E, MS, C, and D) (Extended Data Fig. 10a, b, Fig. 5a, see Methods). To visualize how transcriptional activity is physically arranged, we plotted the average transcript signal for each lineage in Voronoi polygons representative of the spatial arrangement and nuclear space occupied by each region (transcript Gaudi plots, Fig. 5, Extended Data Fig. 11), as done previously^68,69^. Each cluster displayed a different pattern of transcript signal in different lineages (Fig. 5), in line with our prediction that the same higher-order structure can harbour different gene expression patterns that would allow different cell types to carry out different functions. Our analysis revealed that interchromosomal blocks bring together transcriptionally diverse regions of the genome, not exclusively silent or highly active (Fig. 5, green outlines). Instead, spatially contiguous regions of similar activity were more often found outside of interchromosomal blocks. For example, cluster 4 displayed a contiguous stretch of low activity in most lineages, while clusters 5 and 6 displayed stretches of higher activity (Fig. 5). These results indicate that higher-order inter-chromosomal physical proximity is not strictly governed by, or a determinant of, shared transcriptional status (see Discussion).

**Figure 5:**
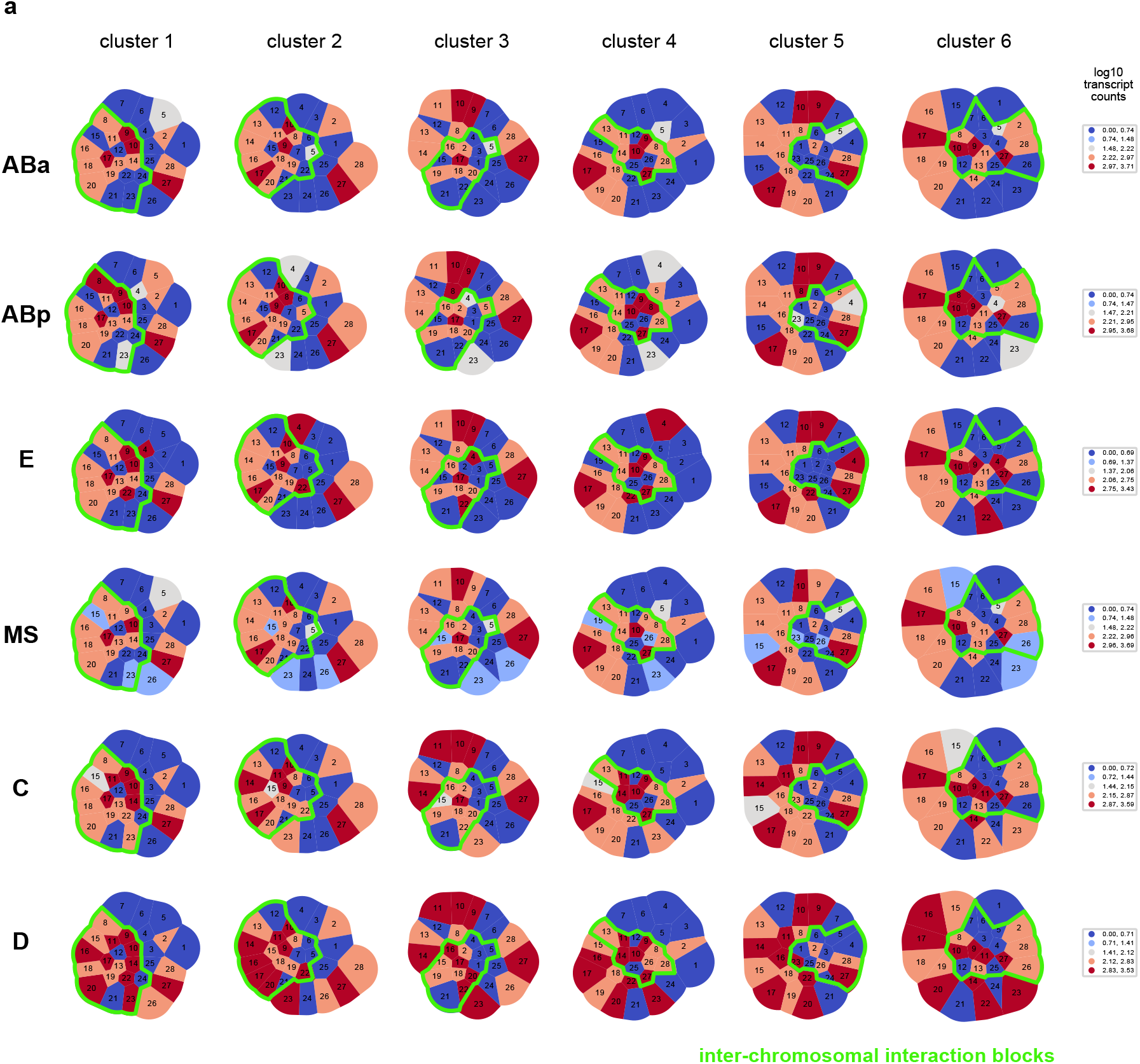
Higher-order interchromosomal blocks coalesce transcriptionally diverse regions. a) Transcript Gaudi plots of chromosome clusters 1-6 in lineages. Each segment represents the space around each chromosome region, and the overall arrangement of segments represents the intra- and inter-chromosomal conformation. Each segment is colored by the total transcript signal in each cell lineage (blue/low to red/high). Segments outlined in green are those present in interchromosomal block associations. Units = log 10 counts.

### 2.6 Local 3D hubs and islands of transcriptional activity differ by structure and lineage

The proportions of higher-order chromosome conformations themselves were not strictly inherited or reproduced between generations within cell lineages, and these global structures did not coordinate loci with similar transcriptional activity when averaged across lineages. We next tested whether the relationship between more local chromatin conformation and transcriptional activity was a stronger heritable feature in embryogenesis. We identified regions that are statistically enriched in a local neighbourhood for transcription as a direct readout of genome function in space, by adapting spatially-informed correlation analysis using Moran’s *I*^70^, which is commonly used in geospatial studies and has recently been applied to the integration of bulk chromosome conformation and ChIP-seq data ^68,69^. Our approach examined the spatial arrangement (from single-molecule chromosome tracing) of gene expression (from scRNA-seq) to categorize genomic regions according to both their transcriptional status, as well as the transcriptional status of neighbouring genomic regions in space. This analysis specifically allowed us to test whether or not transcriptionally active regions clustered in space with other active regions (or not), and whether these local patterns could be inherited within cellular lineages. For each region, we considered its own activity, determined if its local surroundings are enriched for a particular activity level, and classified them into the following 4 categories of ‘metaloci’^69^ (see Methods, Fig. 6a): 1) A region of high transcription surrounded by regions of high transcription (**HH**), representing a 3D hub of high activity. 2) A region of low transcription surrounded by regions of low transcription (**LL**), representing a 3D hub of low activity. 3) A region of high transcription surrounded by regions of low transcription (**HL**). 4) A region of low transcription surrounded by regions of high transcription (**LH**). All regions that are designated in these categories are significantly enriched using local Moran’s *I* with a p-value < 0.05 using a permutation test (Fig. 6a).

**Figure 6:**
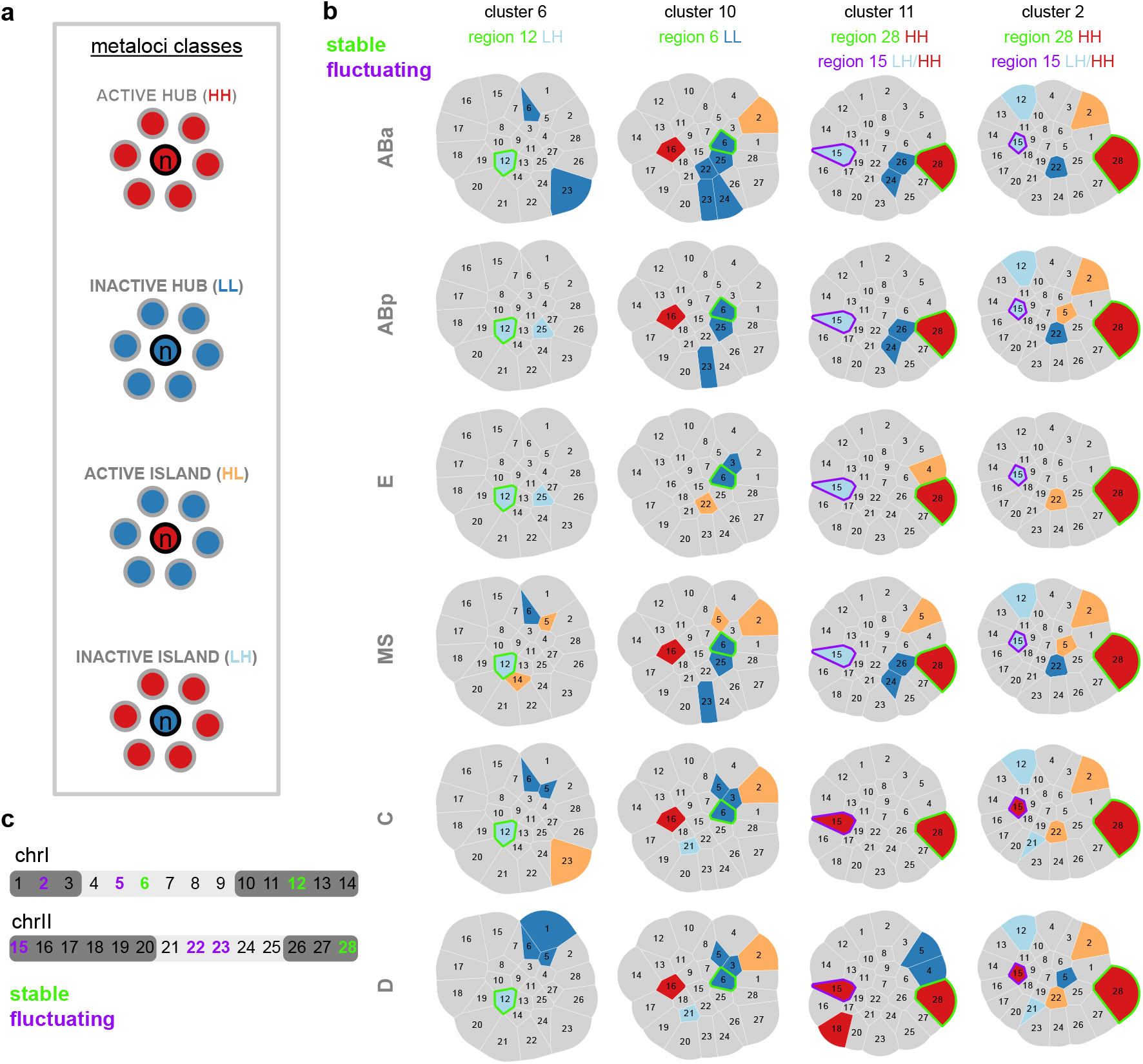
Local 3D hubs and islands of transcriptional activity differ by structure and lineage. a) Schematic of metaloci classes. The dots are coloured blue-red for low-high transcriptional status. The four metaloci classes are hereafter coloured in the following manner: HH regions are coloured red, LL regions are dark blue, HL regions are orange, and LH regions are light blue. b) Metaloci Gaudi plots of example stable and fluctuating regions across lineages. Metaloci classes are coloured as in the labels of Fig. 6a. Regions that maintain a consistent metaloci status across lineages are outlined in green, regions that fluctuate status are outlined in purple. c) Summary of all stable and fluctuating regions across lineages on chrI and chrII schematics.

The prevailing genome compartmentalization model predicts that regions of similar activity will preferentially physically group together into euchromatin and heterochromatin^4^. **HH**/**LL** metaloci are consistent with previous compartment models because they represent spatial hubs of similar activity. On the other hand, **HL**/**LH** metaloci represent regions that resemble islands which maintain their activity in a sea of dissimilar regions, possibly resisting external influence. We found evidence supporting the prevalence of both hubs and islands across structures and lineages in embryos. Metaloci were found distributed across chromosome structures, both inside and outside of interchromosomal blocks (Fig. 6b, Extended Data Fig. 11). Cluster 8 and C lineage structures showed the highest proportions of exclusively **HH**/**LL** hubs, indicating the canonical model of physical grouping of similar regions dominates in these cases. The ABp lineage, E lineage, and cluster 4 showed the highest proportion of exclusively **HL**/**LH** islands, indicating regions are more likely to resist the effects of their surroundings in these cases. However, the vast majority of higher-order structures (48/66) consisted of mixed models containing both hubs and islands (Extended Data Fig. 11). Clusters 1 and 2, which have strong interchromosomal blocks, and the MS lineage contained the highest amount of both hubs and islands. Altogether, these data show that both transcriptional hubs and islands co-exist frequently in the same nucleus at both the intra and interchromosomal level, and support a more nuanced view of chromosome compartmentalization than one based solely on homotypic interactions.

We next asked which specific regions of chrI and chrII showed differential patterns of metaloci classification and distribution across structures and lineages. Some metaloci were consistent across lineages, while others were classified as islands in some contexts but hubs in others. Consistent or stable state regions included: region 6 (chrI centre) **LL** in cluster 10, region 12 (chrI right arm) **LH** in clusters 1 and 6, and region 28 (chrII right arm) **HH** in clusters 2 and 11 (Fig. 6b,c). For region 28, a possible underlying gene causing its consistent **HH** status could be the broadly-expressed translation initiation factor *eif-3*.*B*, which would be required at high levels in all cells for protein production. On the other hand, we identified several regions that are metaloci which toggle states between cell lineages: region 2 (chrI left arm) in cluster 9, region 15 (chrII left arm) in clusters 2 and 11, region 22 (chrII centre) in cluster 2, and region 23 (chrII centre) in cluster 7 (Fig. 6b,c, Extended Data Fig. 11). These fluctuating regions could fall into any class (**HH**/**HL**/**LH**/**LL**). For example, region 15 was **LH** in more anterior lineages (AB, MS, E), but **HH** in more posterior C and D lineages (Fig. 6b). Therefore it appears this region is consistently embedded in a high transcription environment, but is itself silenced in the anterior lineages. There was no overlap between stable regions and fluctuating regions (i.e. stable regions in one cluster did not fluctuate consistently in another and vice versa) (Fig. 6b,c, Extended Data Fig. 11). Additionally, stable regions were not found in the left arms of either chrI or chrII, and fluctuating regions were not found in the right arms of either chromosome. These data indicate metaloci stability is highly influenced by genomic location, chromosome conformation, and lineage identity.

Temporal regulation of gene expression is a crucial feature of embryogenesis, and our metaloci analyses of average lineage transcriptomes in the chromosome clusters indicated that some metaloci have consistent patterns. Therefore, we hypothesized that within each cell lineage, the conformation-transcription relationship may change over time, as both aspects are not static during lineage specification. To test this, we examined metaloci occurrences in single-molecule chromosome traces of individual cells over time in the embryo (see Methods). We split the frequencies of metaloci by class (**HH**/**HL**/**LH**/**LL**) and cell identity (e.g. AB, ABa, ABp, etc.) to produce single-cell conformation-transcription ‘fingerprints’ (Fig. 7, Extended Data Fig. 12). Consistent with the whole-lineage analysis (Fig. 5), the most striking example of a stable region was region 12 (chrI right arm), which occurred as an inactive island embedded in an environment of high transcription (**LH**) in all lineages at high frequency (Fig. 7d, Extended Data Fig. 12). Region 28 (chrII right arm), which was a stable **HH** in cluster 2 and 11 chromosome traces (Fig. 5c), appeared often in this state through embryogenesis, but after the 28-cell stage it was also observed as an **HL** (Fig. 7a,c, Extended Data Fig. 12). These data indicate that the regions physically surrounding region 28 become silent over time but region 28 itself can maintain its high activity. We further identified the stage and cells which were the source of the fluctuating metaloci regions (2, 5, 15, 22, 23) (Fig. 5c-d). For example, region 2 (chrI left arm) occurs most often as an active island in a sea of low transcription (**HL**). However, in certain cells of the E-, C- and D-lineage, region 2 switches strongly to being classified as an **LL** hub (Eal, Ear, Ca, Cp, D), but later regains **HL** status (Extended Data Fig. 12). Similarly, region 15 (chrII left arm) was most often inactive in a sea of high transcription (**LH**) in the AB-, E-, MS-, and C-lineages but not the D-lineage, where it was **HH** in D and Da cells and **HL** in Dp cells (Fig. 7d, Extended Data Fig. 12). Region 2 also occurred as **HH**/**HL** in 4-8-cell AB cells and Cxx cells at lower frequencies (Fig. 7a,c, Extended Data Fig. 12). These data identify regions of the genome that make dramatic and rapid transitions from one transcriptional state and 3D environment to another in different cells.

**Figure 7:**
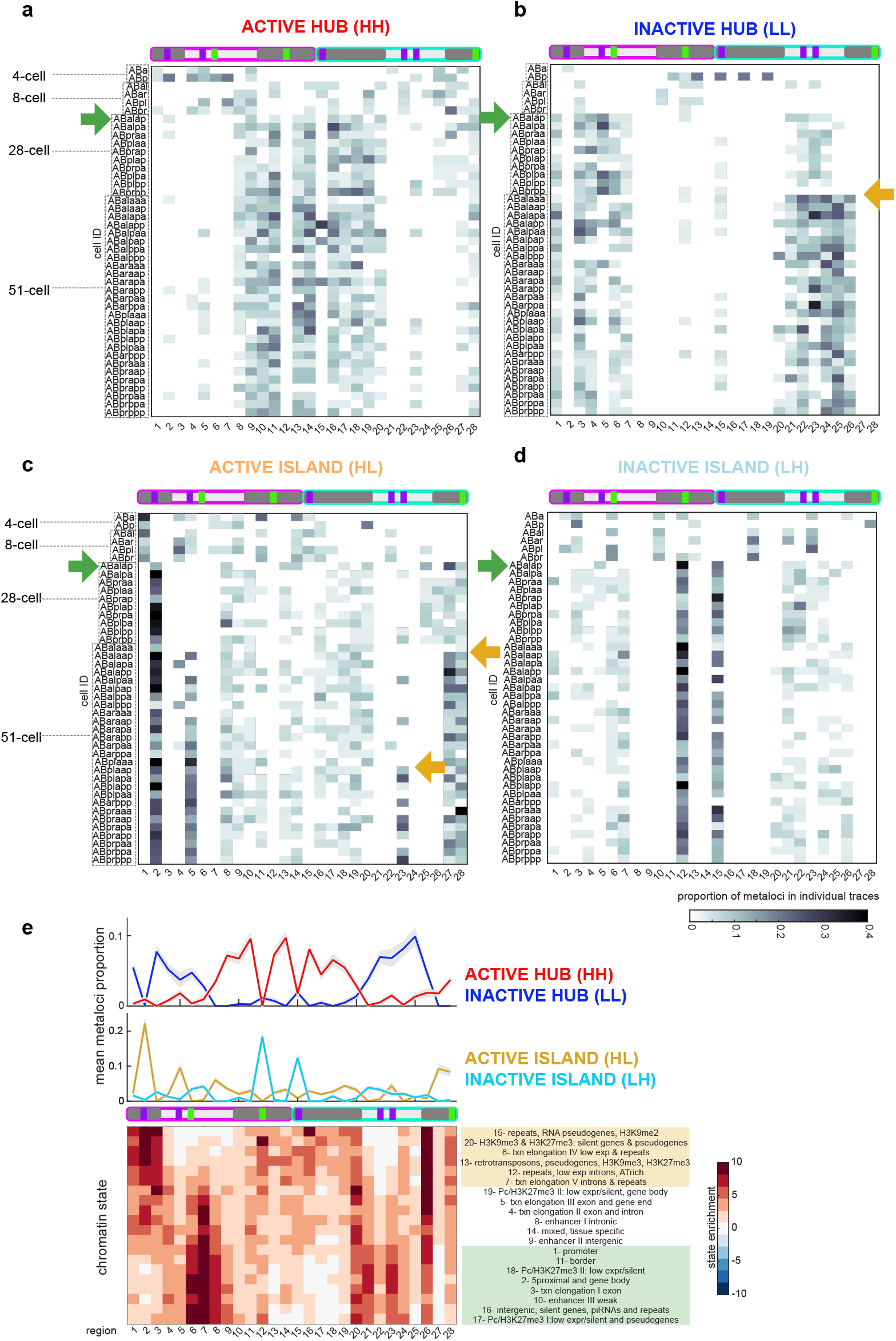
Multiple temporal transitions and canalization of conformation-transcription fingerprints during lineage specification rewire chromatin classes. a) Heatmap representing HH metaloci proportions in chromosome tracing regions of AB-cells over time. b) Heatmap representing LL metaloci proportions in chromosome tracing regions of AB-cells over time. c) Heatmap representing HL metaloci proportions in chromosome tracing regions of AB-cells over time. d) Heatmap representing LH metaloci proportions in chromosome tracing regions of AB-cells over time. e) Chromatin state enrichment^75^ in chromosome tracing bins. Light orange box highlights canonically heterochromatin states, light green box highlights canonically euchromatin states. The mean metaloci percentage signal from AB cells in each category is shown in a line graph above the heatmap, the shaded area represents the standard deviation. Chromosome schematics as in Fig. 4, 6. Green arrows indicate the major ZGA conformation-transcription fingerprint transition. Orange arrows represent more localized transitions.

### 2.7 Multiple transitions and canalization of local conformation-transcription fingerprints during lineage specification

Previous imaging work has shown that the segregation of heterochromatin and euchromatin occurs gradually over *C. elegans* embryogenesis, ultimately culminating in large swathes of electron-dense heterochromatin and H3K9me-positive hubs^71,72^. Bulk biochemical assays have also mapped heterochromatin to broad domains of chromosome arms, and euchromatin to chromosome centres^73–75^. However, due to previous technical limitations, heterochromatin and euchromatin have never been mapped in a lineage-resolved manner with sequence specificity and high temporal resolution. **HH** and **LL** fingerprints present a new way of measuring euchromatin and heterochromatin respectively that remove these limitations. As expected for distinct chromatin types, we find that **HH** and **LL** fingerprints in all lineages appear as complementary and mutually exclusive profiles over genomic locations (Fig. 7a,b, Extended Data Fig. 12). Across lineages, we found that cells in 4-8-cell embryos showed low frequencies of **HH** metaloci in chromosome centres and **LL** metaloci in chromosome arms, similar to the previous definitions of heterochromatin and euchromatin (Fig. 7a,b, Extended Data Fig. 12). The D-lineage is born the latest and has a similar pattern of **HH** centres and **LL** arms at its birth (Extended Data Fig. 12). However, this expected pattern was rapidly eliminated by the 28-cell stage in most cells, and a major transition to a distinct stronger pattern emerged, coincident with the major zygotic genome activation^76^ (see Discussion). In the C-lineage, this transition was also detected in Ca and Cp cells (15-cell stage, Extended Data Fig. 12). The ZGA fingerprints contained high **HH** frequencies predominantly in chrI’s right arm and chrII’s left arm, and a complementary **LL** pattern in the remaining central and arm regions of chrI and chrII (Fig. 7a,b, Extended Data Fig. 12, green arrows). These results suggest that the genomic composition of both heterochromatin or euchromatin are overhauled during the ZGA. Moreover, the major ZGA transition of chrI’s right arm and chrII’s left arm was maintained in subsequent stages, indicating that a stable memory of the conformation-transcription fingerprint persists through cell divisions.

In addition to the major ZGA transition, we also observed more localized transitions occurring in the AB lineage (Fig. 7, orange arrows). For example, some switch-like changes occurred at the 51-cell stage in chrII’s right arm (regions 27 and 28) to increase **HL** frequency, and regions 20-26 to increase **LL** frequency (Fig. 7b,c). Anterior and posterior cells of the AB-lineage bifurcated their fingerprints at region 23, with the posterior cells specifically increasing **HL** frequency (Fig. 7c). All together, these data indicate that chrII’s right arm enriches for overall transcriptional silence over time, with pockets of high activity. Thus, we show that early embryonic stages undergo not only major transcriptional activation in the embryo, but a re-wiring and canalization of chromosome conformation-transcription fingerprints.

### 2.8 Active hubs contain canonically constitutive heterochromatin

Our metaloci analysis suggests that cellular lineages have chromosome conformation-transcription fingerprints that are inherited, and canalized at critical developmental timepoints. To investigate which genes or chromatin classes are enriched in the regions undergoing the major ZGA conformation-transcription transition (Fig. 7a-d, Extended Data Fig. 12, green arrows), we determined the prevalence of 20 previously-defined embryonic chromatin states^75^ along the probed coordinates of chrI and chrII (Fig. 7e). Unexpectedly, this analysis revealed that chrI’s right arm and chrII’s left arm (regions 10-19), which become active hubs (**HH**) at the major ZGA transition, were enriched for canonical heterochromatin classes including pseudogenes, repeats, and retrotransposons (Fig. 7e, orange box), not canonical euchromatin classes (Fig. 7e, green box). Complementary to active hubs, most regions of chrII that become inactive hubs (**LL**), were enriched for canonical euchromatin classes including enhancers, promoters and transcription elongation (Fig. 7e, green box). On the other hand, not all canonically heterochromatin regions in chrI and chrII became active hubs during embryogenesis. For example, region 2 is highly enriched for repeats and pseudogenes, and transitioned to being an active island (**HL**) (Fig. 7c). The left arm of chrI (regions 1, 3) is enriched for heterochromatin classes and predominantly transitioned to **LL**. Region 12 (chrI right arm) is highly enriched for silent genes and pseudogenes with typical heterochromatic histone modifications H3K9me/H3K27me, and transitioned strongly to **LH** (Fig. 7d), indicating that these genes are selectively silenced by the 28-cell stage while being embedded in a 3D environment of high transcription. These data indicate that the genomic makeup and context of these chromatin classes both strongly influence what transcriptional activity a region and its surroundings will adopt during embryogenesis. In contrast to previous population analyses on mixed-stage embryos which have generally defined chromosome arms as heterochromatin-enriched, and chromosome centres as euchromatin-enriched^73,74,77,78^, we now find that the genomic composition of heterochromatin and euchromatin is remodeled during the ZGA transition.

## 3 Discussion

Genome organization regulates transcription, but how different cell types package their DNA *in vivo*, and how this regulation is inherited across cell divisions during embryogenesis are poorly understood. In this study, we developed a new algorithm for cell lineage identification (CeSCALE) combined with single cell genomics to reveal that local conformation-transcription relationships (‘fingerprints’) are associated with, and inherited along cellular lineages. CeSCALE provides a fully automated framework for measuring and quantifying lineage-resolved individual cellular phenotypes in their native biological context, without the need for live imaging, and with near-perfect accuracy for a large developmental window. By combining CeSCALE with single molecule chromosome tracing we uncover prevalent large-scale interchromosomal block associations, which surprisingly coalesce transcriptionally diverse domains and are independent of lineage identity. Instead, we show that local conformation-transcription spatial ‘fingerprints’ are inherited along cellular lineages, by integrating lineage-resolved chromosome conformations with lineage-resolved single-cell transcriptomics. Finally, we find that these ‘fingerprints’ canalize at key developmental stages and represent the rapid re-wiring of chromatin states during the early process of lineage specification.

CeSCALE is a novel computational framework to automatically identify cell lineage identities in *C. elegans* embryos from static snapshots. This framework requires only the 3D centroid positions of nuclei in sample embryos as input, which can be obtained from a variety of image segmentation methods, from manual to deep-learning enabled (see Methods,^79,80^). Our current implementation of CeSCALE uses the P-cell location as an anchor point for the alignment of test and reference nuclei distributions, but future implementations can utilize an arbitrary number of known anchor points depending on the user’s system. Furthermore, CeSCALE per-cell confidence calculations can be used to analyze mutant embryos and identify any cells that deviate from the expected wild-type positions. One limitation of the successful implementation of CeSCALE is that it is dependent on the accurate number of sample nuclei, since test nuclei are matched 1-1 with reference identities. Compared to previous lineage mapping or tracing methods for *C. elegans*^29–33,81^, CeSCALE offers several new advantages: It operates on static images from either live or fixed samples, eliminating the need for temporal information between cells (i.e., tracking cell divisions). CeSCALE does not require manual correction after labeling. It can successfully identify the lineage identities of all embryonic cells in a wide developmental window (4-cell to the 360-cell stage). Finally, the cell identity information gathered from CeSCALE can be applied to any biological question that involves measuring lineage-resolved cellular phenotypes *in situ*, thus providing a broadly valuable resource for the *C. elegans* community.

Embryonic chromosome conformations displayed pervasive territory intermingling. Chromosome territories were originally proposed to form distinct non-overlapping volumes of limited inter-chromosomal interactions in eukaryotic nuclei^64,65,82^. However, notable examples including the intermingling of territories to coalesce ribosomal RNA genes at nucleoli, olfactory receptor enhancers at ‘Greek islands’, nuclear speckles, and telomeres and centromeres in Rabl’s configuration, have shown that interchromosomal contacts have extensive gene regulatory potential^83–87^. Much of the previous work has found that inter-chromosomal contacts were associated either with gene activation or silencing, and in discrete enhancer/gene foci. In contrast, we found that *C. elegans* autosomes displayed unique features where they frequently fold together in large blocks spanning up to half of each chromosome. It is possible that these blocks were not previously uncovered because prior studies focused on tracing single chromosomes^12,13^, or used genome-wide bulk measurements of contact frequency that favor intra-chromosomal interactions and dilute possible cell-cell differences by averaging^58–62^. It will be interesting in the future to test whether large interchromosomal blocks are conserved in other model systems of embryogenesis.

Surprisingly, interchromosomal blocks contained not exclusively silent or active domains, but a mix of both in lineage-specific patterns. This result has implications for our current models of genome compartmentalization, which propose that compartments bring together distal regions of similar epigenetic status or transcriptional activity in space^4,5^. At the kilobase level, recent studies have indeed shown that fine-scale compartments correlate well with uniform epigenetic and transcriptional status^59,62,88,89^. In contrast, our data show that transcriptionally diverse domains fold together often at the Megabase level, arguing against activity being an absolute mechanistic driver of physical proximity and vice versa. Alternatively, the large interchromosomal blocks detected in this study may contain smaller scale, more intricately organized sub-compartments which correlate more positively with uniform transcriptional status. However, in line with our observations, metaloci analysis revealed that particular regions of the genome can maintain their activity status despite being surrounded by dissimilar chromatin (LH and HL classes). LH/HL are chromatin classes that could not be previously predicted based solely on histone modifications or transcriptional activity - they were uncovered by virtue of our cell identity-resolved metaloci analysis, and may represent novel chromatin subtypes with unique properties that allow them to resist the influence of surrounding repressive or activating factors.

Local chromosome conformation-transcription relationships canalized during early embryogenesis. Leveraging lineage relationships, we determined that conformation-transcription fingerprints, not conformations alone, are inherited through cell divisions. Previous studies have characterized the heterochromatic and euchromatic elements of the *C. elegans* genome at the population level in embryos of mixed-stages^73,74^, while in our study we define chromatin classes with high spatial and temporal resolution. HH and LL metaloci classes represent euchromatin and heterochromatin respectively^69^, based on transcriptional activity and chromosome conformation. We found that HH and LL have complementary frequencies across chromosome regions, as expected from opposing chromatin types, but surprisingly lose the expected genomic locations based on previous ChIP-seq analyses^73,74^ at the 28-cell stage in most cells. This stage corresponds to the major wave of transcription that occurs during the zygotic genome activation^76^, suggesting that chromosome conformation and transcription undergo large transitions concurrently. Thus, our data indicate that previously unknown shifts occur in the genomic compositions of heterochromatin and euchromatin at the single-cell level throughout embryogenesis.

This work provides CeSCALE as a new valuable resource for cell lineage analysis. Moreover, integrating CeSCALE with single-cell genomics highlights that each cell during early development carries a fingerprint of its chromosome conformation-transcription relationship, suggesting chromosome conformation itself is fluid enough to facilitate or adapt to the rapidly changing transcriptome.

## 4 Methods

### 4.1 Cell alignment problem formulation

In order to align the cell IDs/labels from a *reference* embryo image (labeled), denoted as k, to a *test* embryo image (unlabeled), denoted as k’, we will compare the two corresponding distance matrices (i.e., all-by-all cell distances) and try to rearrange the one of them such that they become as similar as possible (e.g., in the Euclidean sense). After doing so, we can get the cell IDs correspondence by the aligned rows/columns. The mapping from the labeled cells to the unlabeled ones can be formally described by a *permutation matrix ↟* of size *n* × *n* (where *n* is the number of cells in the embryo image) such that 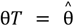, where θ is the vector containing the cell IDs of the labeled image (reference) and 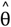 is the vector with the assigned names for the unlabeled image (test). Thus, inspired by^36^, we formulate the cell alignment problem as a quadratic assignment problem^90^ where we aim at finding the optimal permutation matrix *T*^*^ that minimizes the discrepancy between the two distance matrices (i.e., reference and test distance matrices):

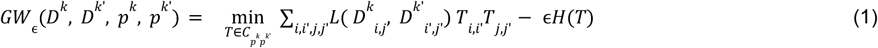

where *L*(·) is the loss function and here we select it as the quadratic loss 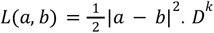 and *D*^*k*^ are the reference and test distance matrices, respectively. *p*^*k*^, *p*^*k*’^ are probability measures (here assumed to be uniform), which can be seen as prior weights for every cell appearing in embryos k and k’, respectively, ϵ≥0 is the regularization term and

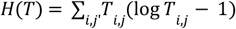

is the entropic regularization term. *T* is a coupling between the two spaces on which *D*^*k*^ and *D*^*k*’^ are defined and

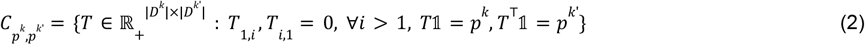

The reason why above we set all elements of the resulting transportation plan in the first row and column equal to 0 has to do with the fact that the P-cell is known and will be explained in detail in the next section. Finally, we note that in Eq. (1) when the regularization term ϵ above is small then *T* is close to a (scaled) permutation matrix and when it is large then *T* is *fuzzy* (i.e., has many non-zero entries). As a result, higher values of ϵ promote more fuzzy transportation plans.

### 4.2 Extending the algorithm to incorporate information about the P-cell

For the problem under consideration, we note that since the identity of the P-cell is known, this information should be incorporated into the optimization problem. We make the convention that the position of the P-cell is always the first position in the distance matrices, meaning that the first row and column of all distance matrices encode the distance of the respective P-cell with all other cells in the image. Then, we can add the following additional constraints in the problem (*see* (2)).

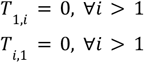

We can take advantage of the fact that the elements of the first row and column of the transportation matrix are all zero except for the element *T*_1,1_ and explicitly incorporate the above constraints into the optimization problem by simplifying the summation in the objective function (Eq. (1)) as follows.

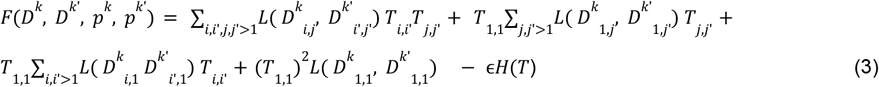

We note here that *T*_1,1_ is not an optimization variable and has a fixed value. Specifically, for the case of uniform *p*^*k*^ and *p*^*k*’^ and |*D*^*k*^| = |*D*^*k*’^| = *n*, we have *T*_1,1_ = 1/*n*. Taking the derivative w.r.t. *T*_*i,i*_ for *i, i*’ > 1 and using the fact that the distance matrices are symmetric (i.e., *D*_*i,j*_ = *D*_*j,i*_ for all *i, j* ∈ {1, ..., *n*}), we get

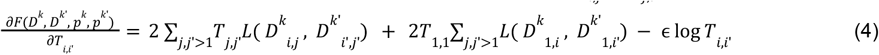

Thus, the gradient ∇_T_F is given by a *n*-by-*n* matrix where the first row and column are zeros and the rest of the entries are given by Eq. (4).

This concludes the derivation of the extension of the algorithm presented in^36^ to and we solve the optimization problem (1) using projected gradient descent as in^36^ while we use the derivatives computed in Eq. (4) to obtain the submatrix. *T*_2:n,2:n_ (since the first row and column of *T* matrix are fixed in our case)

We note that the obtained solution is not independent of the entries of first row and column of *L*( *D*^*k*^, *D*^*k*’^), since the gradient (Eq. (4)) includes information about the first row and column through the term 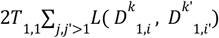

### 4.3 The end-to-end pipeline

After presenting the modified GW computation, which is the base of our algorithm, we will present in the sequel the end-to-end process we follow for cell-lineage identification from static images.

Let us assume we have access to a set of reference embryos (*reference cohort*), where the cell IDs are known, and a test embryo with unknown cell IDs (except for the P-cell which is marked and known). All embryos are of the same age *n* and for all of them we have access to their cell coordinates on the X,Y,Z axes. All embryos in the reference cohort share the same set of cell IDs (“patterns”). If more than one pattern is available for a given developmental stage, then the algorithm evaluates all candidate patterns and assigns to the test embryo the pattern with the best fit as explained in more detail in step 5 below. The algorithmic steps are the following:

1. **Axes scaling and distance matrix computation.** We use the Euclidean distance metric to create the distance matrix (DM) for each embryo, where each entry *i, j* represents the distance between the cells corresponding to row *i* and column *j*. The matrix is symmetric with zeros on the diagonal. After applying the algorithm on the original distance matrices, we observed that beyond developmental age 28, errors began to accumulate. Upon closer inspection, we found that these errors stemmed from the disproportionate influence of the axes: because the z-axis was highly condensed, its contribution to the Euclidean distance was relatively small. As a result, many cells that were spatially distinct along the z-axis—but close in the x–y plane—were misclassified, often leading to label flips. For this reason, we applied Z-axis upscaling, thereby increasing the relative weight of the z-axis values in the distance computation. Working with distance matrices computed on this transformed space significantly increased accuracy. Empirically, we found that the most effective scaling ratio was approximately 1:1:0.8 for axes X,Y,Z, corresponding to physical scales 254nm, 254nm, 200nm, respectively. In practice, since the initial labeled data used axis scales of 254nm, 254nm, 1000nm for X,Y,Z axes, respectively, we achieved this effective ratio by multiplying the Z-axis values by a factor of 5.
2. **Reference maps.** We create the reference maps by computing the average (arithmetic mean) of the distance matrices of the embryos in the reference cohort in order to get a more robust estimate of the average cell distances.
3. **Cell alignment of the test image.** This step constitutes the most vital step of the process. Here we obtain the cell labels for the test embryo. The procedure and mathematical details are explained in detail in Sections 4.1, 4.2. Before applying the algorithm, both the reference and test DMs are normalized by dividing with the respective maximum element of each matrix. For the choice of the regularization parameter ϵ, a grid search is performed in the intervals [0.0001 - 0.0009] with step 0.0001 and [0.001 - 0.009] with step 0.001, and the value of ϵ that yields the smallest Gromov-Wasserstein distance (see Eq. (1)) is selected. In this way, we obtain the optimal transportation plan *T*^⋆^ from the Sinkhorn algorithm that indicates how we should re-arrange the rows and columns of the reference DM to match as closely as possible the test DM. However, *T*^⋆^ is not a permutation matrix yet (i.e., *T*^⋆^ contains fuzzy entries or it may have the largest value of two different rows/columns in the same column/row), which we would need so that we perform the 1-to-1 matching of the cells from the reference image to the test image. For this reason, we introduce the next step.
4. **Maximum weight full matching.** In order to obtain a permutation matrix from the transportation plan *T*^⋆^ is to select the edges that maximize the overall sum. This is the *maximum weight full matching* problem for bipartite graphs, which can be expressed as a *linear program*, and solved by^91^. In this way, we obtain the permutation matrix *W*^⋆^, which indicates the assignment of the cell IDs to the test embryo. More specifically, let θ be the vector of the cell IDs of the reference embryo, we obtain the assigned IDs for the test embryo as θ*W*^*^.
5. **Embryo super-impositioning.** After we obtain the cell labels for the test embryo image, we can perform *super-impositioning* of the test embryo onto the reference cell positions in order to check the spatial agreement of the assigned cell IDs and whether the assigned cell IDs of the test embryo are close to the expected cell positions or not. To do so, we need to transform the cell distances of the reference DM into coordinates in space. We call these coordinates as *reference coordinates* and we obtain them by applying Multi-Dimensional Scaling^92^ to the reference DM. After doing so, we super-impose the assigned Cell IDs of the test embryo onto the reference coordinates by applying the Procrustes algorithm in order to find the best translation, rotation and scaling so that the test Cell IDs coordinates match as well as possible the reference coordinates^48,49^.This provides a way to visually evaluate the outcome of the alignment, as well as a way to compute a quantitative measure of spatial proximity of the alignment, the *normalized Procrustes distance* between the reference cell positions and the respective aligned cells with the same ID in the test image. The normalized Procrustes distance is given by:

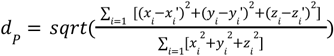

where *x*_*i*_, *y*_*i*_, *z*_*i*_ are the X,Y,Z-axis reference coordinates of cell i and *x*_*i*_’, y_i_’, *z*_*i*_’ are the aligned (after Procrustes alignment) coordinates of the test embryo. *d*_*P*_ takes values between 0 and 1 and the smaller this distance is, the better the alignment of the test embryo with the reference coordinates is. Due to the known slight variability in the timing of some cell divisions in the embryo^93^, we note here that for some stages there can be only 1 possible set of cell IDs (patterns), while for some others there can be more than one pattern. We call the stages belonging to the first category unimodal, while the latter ones multimodal. In case we deal with a multimodal stage, then we select the pattern with the smallest Procrustes distance, as this indicates better cell alignment for that set of cell IDs. A list of the stages along with their corresponding patterns available in the labeled data we had access to can be found in the Supplementary Data 1.
6. **Cell-confidence scores computation.** The normalized Procrustes distance provides a good measure of global alignment, but to assess *local alignment accuracy* at the single-cell level we introduce the so-called *cell confidence scores*. Specifically:
  a. Within the reference cohort, we first perform Procrustes alignment across all images to align the coordinates of the Cell IDs of the embryos in the reference cohort.
  b. For each cell ID, we then apply Kernel Density Estimation (KDE) (with Gaussian kernels) to model the probability distribution of its 3D spatial positions.
  c. For every cell ID of the test image, after the alignment with the reference coordinates (step 5), we compute this per-cell confidence score as the likelihood ratio of the observed cell position under the KDE distribution of the corresponding reference cell ID. Formally, for a cell i and a cell ID j, the confidence score that i corresponds to cell ID j is computed as:

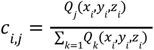

where *Q*_*j*_ (*x*_*i*_, *y*_*i*_, *z*_*i*_) is the probability density (as estimated via KDE) for cell ID j at position (*x*_*i*_, *y*_*i*_, *z*_*i*_). The denominator normalizes over all possible cell IDs *k*, ensuring that: ∑_*j*_ *c*_*i,j*_ = 1, for all *i*. Thus, *c*_*i*_ represents the posterior confidence that cell *i* corresponds to identity *j*, given its spatial location relative to the reference cohort and a high confidence score indicates that the predicted cell position falls within the expected spatial distribution of that cell ID in the reference cohort, whereas low scores reflect atypical placements. For this step, larger reference cohorts give better results, as the probability distribution modelled by KDE is more consistent.

One important remark here needs to be mentioned. When applying the algorithm to a test dataset that has different scales, one extra preprocessing step should be performed before the 6 aforementioned steps, in order to normalize the axis scales between reference and test data. More specifically, the test axes should be scaled back to the reference scale (e.g., here the scale is 254nm, 254nm, 1000nm for X,Y,Z axis, respectively) and then follow the six steps of the process shown above. An example of applying CeSCALE to data of different axis scales is presented in Section 4.5 in the sequel.

### 4.4 Accuracy computation of CeSCALE on live imaging data

Let’s now describe in detail how the accuracy scores of CeSCALE were computed. To evaluate accuracy, we used published annotated live-imaging data from 48 wild-type *C. elegans* embryos spanning the 2-cell to 360-cell stages, with frames captured every 75 seconds (Fig. 1a,^45–47^), yielding 3923 images in total. Cell identities and nuclear centroids (X, Y, Z coordinates) were extracted using Acetree/StarryNite followed by manual curation (Fig. 1b,^45–47^). Because only certain developmental ages were sufficiently represented, we set a minimum threshold of 40 images per age to be included in the analysis. The list with the stages and cell patterns for every stage with sufficient data can be found in Supplementary Data 2.

For each developmental age, we computed the accuracy of CeSCALE for every available image as follows. First, we applied step 1 of the algorithm (see Section 4.3) to all images. Then, in step 2, for each image we generated a reference map by averaging the distance matrices of all other embryos at the same age, in a 1-vs-all design. Importantly, images from the same embryo as the test image were excluded from the reference cohort which the reference map is generated from. This ensured that the reference map did not inadvertently include images from the same embryo; although successive images are not identical due to cellular movement, their similarity could otherwise introduce data leakage through temporal correlations. Steps 3 and 4 were then applied to the test image to assign cell labels. Finally, the predicted labels were compared to the ground truth to measure algorithmic accuracy. The average accuracy of the image is given by 1-N_errors_/N_total_, where N_errors_ denotes the number of incorrectly labeled cells (i.e., wrong cell ID assigned to them) and N_total_ corresponds to the total number of cells. Thus, a value of 1 is equivalent to 100% accuracy.

Applying this methodology across all developmental ages and all available images yielded the accuracy results summarized in Extended Data Fig. 2a, where we observe that the average accuracy is remarkably high. One notable exception occurred at the 4-cell stage where accuracy is significantly lower. This reduction is explained by the symmetry of the 4-cell embryo: the relative distances between ABp and EMS to other cells are highly similar, making them difficult to distinguish. At later stages, as this symmetry breaks, the accuracy rises sharply. Cell-specific performance is illustrated in the dendrogram Fig. 2a, which highlights consistently high accuracy across all available cells. The lowest accuracy again occurs for ABp and EMS at the 4-cell stage, consistent with the symmetry-related ambiguity described above.

#### 4.4.1 Procrustes Distance per Embryo

After obtaining the cell identities, for every test image, we superimpose its coordinates onto the respective reference coordinates (for that specific age) and compute the (normalized) Procrustes distance–a measure of the residual shape discrepancy after optimal translation, rotation, and scaling (step 5). We observe in Extended Data Fig. 2b (top figure) that the distribution of these distances is close to zero (average Procrustes distance across all images is 0.02), thus indicating good spatial alignment. Finally, the individual cell confidence values for the annotated data (see Supplementary Data 4) indicate robust cell labelling.

### 4.5 Applying CeSCALE to unlabeled data

Before applying **CeSCALE** to a new dataset, the spatial scales of the X,Y,Z axes must be normalized. The algorithm was originally parameterized and tested for its accuracy on the reference voxel dimensions of [*S*_*x*_, *S*_*y*_, *S*_*z*_] = [254nm, 254n*m*, 1000*nm*]. Therefore, prior to running the algorithm, the coordinates of each embryo in the new dataset must be rescaled to this reference system.

Let *S* = [*S*_*x*_, *S*_*y*_, *S*_*z*_] = [254nm, 254n*m*, 1000*nm*] (reference scales) and *S*’ = [*S*’_*x*_, *S*’_*y*_, *S*’_*z*_] (scales of the new dataset). For each axis, the rescaling factor is given by

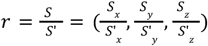

In other words, each coordinate of the new dataset is divided by the corresponding axis-specific ratio to bring all data into the same metric space as the reference cohort.

After this preprocessing step, the algorithm steps are applied as described in Section 4.3. We applied this procedure for the unlabeled data we gathered from fixed embryos (76 embryos in total). As a reference cohort, the entire labeled dataset was used. We note that in order for the methodology to be applicable, the age of the test embryo should be included in the set of available ages in the reference cohort. The full list of available ages and cell patterns in the reference cohort (annotated data) can be found in Supplementary Data 1. For the data gathered for these 76 embryos all ages were present in the set of ages in the reference cohort.

Since for these data we do not have access to the ground-truth labels, we resort to the (normalized) Procrustes distance (step 5) and to the cell-confidence scores (step 6) to investigate the quality of the assigned cell IDs. The Procrustes distance for every image of this dataset is shown in Extended Data Fig. 2b (bottom figure), where the low distance values (average score across all images is 0.06) indicate a robust cell alignment. An example of KDE application (step 6) is shown in Fig. 2b. The average cell confidence per age is depicted in Extended Data Fig. 2c (average score across all images and all cells available is 0.84). See also Supplementary Data 4 and 5 for individual cell confidence values. Supplementary Data 4 shows the individual cell confidence values calculated on the live imaging data (with ground truth labels). Supplementary Data 5 shows the individual cell confidence values for fixed imaging data.

### 4.6 scRNA-seq processing

scRNA-seq data was downloaded from Cole *et al*.^52^, as this study covered the same developmental window as our chromosome tracing results (See Supplementary Data 12 and 13). To remove the signal corresponding to maternally-deposited RNAs and only consider newly generated RNAs during ZGA, we subtracted from each cell the signal for each gene found in the 1-cell embryo. We next binned all transcript data for chrI and chrII to assign each gene’s signal to the closest location of the 28 regions probed by chromosome tracing. After the removal of the maternal contribution, the transcript counts were log-transformed (log base 10). This data was then used for metaloci analysis following Mota-Gomez *et al*.^69^.

### 4.7 Spatial auto-correlation using Moran’s *I*

#### 4.7.1. Cluster-based metaloci analysis

For the results in this section, we utilized 62 embryos, out of which we kept the 52 which had Procrustes distance less than 0.15. Out of these 52 embryos we obtained 4248 chromosomes. Each chromosome is divided into 28 regions, but not all of them are detected. For this reason, we choose to discard the chromosomes that have less than 10 regions identified, resulting in the final amount of available chromosomes to 2975 chromosomes.

To reveal the conformation-transcript relationship between the Clusters and lineages, we applied the Mota-Gomez *et al*.^69^ methodology, with the Cluster median distance matrices being used as well as the average transcript levels (maternal-removed and log-transformed) from the available cells (as identified by CeSCALE) for the following lineages ABa, ABp, MS, C, E, D. The methodology proposed in Mota-Gomez *et al*.^69^ was applied in our context as follows. A graph drawing algorithm (Kawai-Kamada) is applied to the distance matrices and then the Voronoi polygons are constructed. Finally, the Moran I is calculated for the chromosome regions (28 in total) and by performing a permutation test, the p-values are calculated to define which regions constitute metaloci for every lineage-cluster combination (Fig. 6b, Extended Data Fig. 11).

#### 4.7.2 Cell-based metaloci analysis

To further proceed with our analysis on a more granular level (cell-specific), we performed the metaloci analysis on specific cell IDs. We collected the transcript levels of the cells listed in Supplementary Data 12. For these cells, we obtained 2181 chromosomes. Since, we know that chromosome conformations are highly variable, we tried to use cells that were close to the reference age it usually belongs to and we kept only the chromosomes that belong to that reference age within a window of 11 ages difference. This reduced the available chromosomes to 1976. So, for every cell that falls within the aforementioned age-window we used its chromosome distance matrix and its transcript level^52^ to compute the metaloci. Finally, the percentages for each type of metaloci (HH, LL, HL LH) for every cell ID were derived (Fig. 7, Extended Data Fig. 12).

### 4.8 *C. elegans* Strains

Live imaging data was previously collected from N2(wild-type) embryos^46,47,94^. Fixed imaging data in this study was collected from JH3269, which has PGL-1::GFP tagged at the endogenous locus in an N2 background. JH3269 was cultured consistently at 20°C using standard *C. elegans* husbandry^95^.

### 4.9 RNA smFISH

RNA smFISH was performed as previously described^96^, with some modifications. smFISH probes were pre-annealed by adding 4ul primary oligos (100 uM), 1 ul FLAPY/Z (100 uM), 2 ul NEB buffer 3, 13 ul RNase free water. ceh-51 primary probes were annealed with FLAPY-ATTO565 and tbx-37 primary probes were annealed with FLAPZ-ATTO647N. Probe sequences can be found in Supplementary Data 10.

Embryos were fixed in 1xPBS/0.05%Triton/1%PFA at RT for 5 min and adhered to poly-L-lysine coated slides (Epredia™ Epoxy Diagnostic Slides, 10106260). Slides were frozen on a dry ice-cooled metal block for 30min, and coverslips were flipped off with a blade to remove eggshells. Samples were immediately placed in ice-cold methanol for 5min, then washed 4x for 15min each (once in PBS, twice in PBS/0.5%Triton, once more in PBS). Samples were pre-hybridized with smFISH hybridization buffer (2xSSC/ 10% formamide/ 0.1g dextran) at 37C for 1h. Probes were pre-annealed in smFISH hybridization buffer at 1:50 dilution to the final concentration of 3.2 nM for each single oligos, and with GFP-booster ATTO488 (ChromoTek gba488) at 1:200 dilution. Samples were incubated overnight at 37C, then washed with 2xSSC/10% formamide 5min each 3 times. A long wash in 2xSSC/10% formamide 1 h at 37C was performed, followed by two final washes in 2xSSC/10% formamide 5 min each. Samples were stained with DAPI 1:1000 dilution in 2xSSC, then mounted and sealed with Antifade VECTASHIELD®(Vector Labs, H-1000-10). Imaging was done using an Olympus IX83 and Yokogawa CSU-W1 confocal (50 um disk), with an 100x/ NA 1.5 oil UPL APO objective in the Imaging Core Facility of the Biozentrum.

### 4.10 Chromosome tracing by multiplexed sequential FISH

Chromosome tracing was performed as in Sawh *et al*. 2020^12,54^ with slight modifications to the hybridization protocol to include immunostaining. Briefly, fluorescent primary probes coat the genome sequence, and fluorescent secondary probes bind to unique tails on primary probes (specific to each region). Regions are detected in sequential hybridization rounds on the same sample.

Primary DNA FISH probes were designed to twenty eight 100kb regions along the lengths of chrI and chrII using the following structure: 5’ – 20nt PCR-priming sequence – 30nt genome homologous sequence – 30nt readout sequence (to bind to its corresponding unique secondary probe) – 20nt 30 PCR-priming sequence – 3’. A density of 10 individual probes per kb genomic sequence was used, therefore each region was targeted by 1000 tiled fluorescent probes. Homologous sequences were designed using OligoArray2.1 and OligoArrayAux^97^ with the following parameters: melting temperature 60-100°C, no cross-hybridization or predicted secondary structure with a melting temperature greater than 70°C, GC content 30%–90%, no stretches of 7 or more identical nucleotides. Specificity of homologous sequences were further verified using NCBI BLAST+ 2.7.1. PCR priming and readout sequences contained no significant homology to the *C. elegans* genome and were previously validated12. Sequences for all probes can be found in Supplementary Data 9. Probe libraries were synthesized by CustomArray (now GenScript), amplified and conjugated to fluorophores as previously described^12^. ChrI primary probes were 5’ labeled with ATTO 565 and chrII primary probes were 5’ labeled with AlexaFluor647 (IDT).

Hybridization of DNA FISH probes to embryos was performed as previously described^12,54^, except after fixation and before DNA FISH, embryos were immunostained. Samples were blocked with 5%BSA in PBS for 1h at room temperature then incubated with a mouse anti-GFP antibody (MilliporeSigma MAB3580, 1:500 dilution in 5%BSA) overnight at 16°C to detect the P-granule marker (PGL-1::GFP). Three washes were done in PBS/0.5% Triton for 15min, followed by secondary antibody incubation (Goat anti-mouse IgG (H+L) AlexaFluor 488, Fisher Scientific A11029), and three additional washes. Samples were then fixed with 1%PFA/PBS/0.05%TritonX100 for 10 min before proceeding with DNA FISH using the primary chromosome tracing probes.

Imaging was performed as previously described using a widefield fluorescence microscope (Nikon Ti2E) coupled to a microfluidics system to deliver secondary probes to the sample and perform washes in sequential rounds^12,54^. Our custom code to control the microscope and fluidics system can be found on GitHub (https://github.com/hilsawh/microscope_control). We used a Prime-95B CMOS camera (Teledyne Photometrics), and a 100X Plan Apochromat objective (Nikon). Between each round, probe signal from the previous round was eliminated using photobleaching. Chromatic aberration was calculated for the imaging channels at the beginning or end of each experiment using four-colour beads (Tetraspeck microspheres, Fisher Scientific T7279). Samples were mounted with DAPI (405nm) and fiducial beads (FluoSpheres™ Carboxylate-Modified Microspheres 0.1 μm Yellow-Green 505/515 nm, ThermoFisher F8803) on the coverslip to account for any sample drift between imaging rounds. The first round of imaging captured nuclear volumes, P-granules, chrI and chrII territory volumes, and fiducial beads. Subsequent rounds captured individual chromosome regions (secondary probe signal), and fiducial beads.

### 4.11 Chromosome Tracing Image Analyses

Image analyses were performed in MATLAB (v2023b) as previously described^12^ with modifications to 3D segmentation to identify the P-cell’s nucleus. Our custom code to perform image analysis can be found on GitHub (https://github.com/hilsawh/chromosome_tracing). To define nuclear and chromosome territory volumes, images were thresholded, denoised, binarized, distance transformed, and segmented using marker-controlled watershed segmentation. To define the P-nucleus, P-granule volumes were defined by thresholding and binarization. The centre of mass of all P-granules was calculated (P-granules have perinuclear localization) and the closest nucleus was found by Euclidean distance. This nucleus was then defined at the P-cell nucleus, and all other nuclei were classified as somatic. CeSCALE was then applied to all embryos to label somatic cell IDs.

Construction of chromosome traces was performed as previously described^12^ without modifications. Briefly, foci from secondary hybridization images (regions 1-28) were pinpointed using 3D Gaussian fitting. Next, within each chromosome territory, foci were connected to each other in ascending order using the nearest-neighbour method (i.e. 1 followed by the closest 2, then closest 3, etc.). Chromosome traces were categorized to cell IDs after CeSCALE by finding which nuclei they belonged inside. All traces annotated by cell ID can be found in Supplementary Data 11.

### 4.12 Clustering

Unsupervised clustering was performed using Seurat^98^ 5.0.3 (RStudio, 2023.12.1+402) as previously described^12^ without modifications. We used all pairwise distances between the 28 regions of chrI and chrII to cluster, and a minimum trace length of 10 pairwise distance pairs. Clustering resolution was initially performed at a range of values (0.5-0.9), then the two largest clusters were further subset to obtain clusters that were visually distinct by median pairwise distance matrices and overall low standard deviation. Marker regions were defined in Seurat by Wilcoxon Rank-Sum test and clusters were further compared individually by Welch’s t-test (Bonferroni-corrected).

### 4.13 Chromatin state analysis

Canonical heterochromatin (arm) and euchromatin (centre) compartments were defined as previously described in the literature using ChIP-seq for histone modifications^73,74^. Twenty previously defined embryonic chromatin state maps based on combinations of histone modifications, gene family enrichments, and other genomic features (e.g. enhancers, promoters) were downloaded from Evans *et al*. ^75^. We next binned all state data for chrI and chrII to assign each state’s signal to the closest location of the 28 regions probed by chromosome tracing. The above data were collected from mixed stages of embryos in bulk biochemical assays.

### 4.14 Statistics and reproducibility

No statistical method was used to predetermine sample size and data was excluded upon outlier analysis. If multiple tests were carried out on the same data, p-values were corrected for multiple testing using Bonferroni correction or as stated in the results or methods.

## Supplementary information

Supplementary Data 1. List of stages and corresponding cell patterns along with number of embryos and images available for each in the annotated live imaging data.

Supplementary Data 2. List of stages and corresponding cell patterns along with number of embryos and images available for stages and cell patterns with more than 40 images.

Supplementary Data 3. HTML file for data exploration of live imaging data accuracy.

Supplementary Data 4. HTML file for data exploration of live imaging data confidence.

Supplementary Data 5. HTML file for data exploration of fixed imaging data confidence.

Supplementary Data 6. HTML file for data exploration of 8-cell KDE.

Supplementary Data 7. HTML file for data exploration of 15-cell KDE.

Supplementary Data 8. HTML file for data exploration of 24-cell KDE.

Supplementary Data 9. Probe locations and sequences for chromosome tracing.

Supplementary Data 10. RNA smFISH probe sequences.

Supplementary Data 11. Chromosome traces annotated by cell id.

Supplementary Data 12. Correspondence file between chromosome traces and scRNA-seq cell IDs used for metaloci analysis.

Supplementary Data 13. Reference ages for scRNA-seq cell IDs used for metaloci analysis.

## 6 Acknowledgements

We are grateful to Anthony Santella and Zhirong Bao for generously curating the lineage-annotated live imaging data we used to create our reference maps. We also thank Zhirong Bao and Maxwell Shafer for comments on the manuscript. *C. elegans* strains were provided by the CGC, which is funded by NIH Office of Research Infrastructure Programs (P40 OD010440). RNA smFISH was performed with the support of the Imaging Core Facility of the Biozentrum. Calculations were performed at sciCORE (http://scicore.unibas.ch/) scientific computing center at University of Basel. A.N.S. was supported by the Natural Sciences and Engineering Research Council of Canada (NSERC, RGPIN-2024-04767), the Canadian Institutes of Health Research (CIHR, PJT-197921), a Canada Research Chair in 4D Genomics (CRC, 2023-00193), the Maud Menten Prize in Genetics (PJJ-204167), and the Swiss National Science Foundation (SNF, SPARK CRSK-3_195955).

## 7 Data availability

Lineage-labeled live imaging data (ground truths) were curated from previous publications^45–47^. Chromosome tracing raw image data is available upon request due to the large dataset sizes (TBs) from multiplexed FISH.

## 8 Materials and Code availability

Code generated in this study will be freely available to the public in the following GitHub repositories: https://gitlab.com/ceda-unibas/cell-lineage-identification https://gitlab.com/ceda-unibas/cell-lineage-metaloci-computation https://github.com/hilsawh/microscope_control https://github.com/hilsawh/chromosome_tracing

## 9 Author contributions

K.N.: Conceptualization, Methodology, Software, Validation, Formal Analysis, Investigation, Data Curation, Writing - Original Draft, Writing - Review & Editing, Visualization.

F.X.: Investigation (RNA smFISH).

N.B.: Formal Analysis (Image Segmentation).

G.F.: Supervision, Project Administration, Writing - Review & Editing.

H.P.M.: Methodology.

I.D.: Supervision, Project Administration, Writing - Review & Editing.

S.E.M.: Conceptualization, Resources, Writing - Review & Editing, Supervision, Project Administration, Funding Acquisition.

A.N.S: Conceptualization, Methodology, Software, Validation, Formal Analysis, Investigation, Data Curation, Writing - Original Draft, Writing - Review & Editing, Visualization, Supervision, Project Administration, Funding Acquisition.

---

**Extended Data Fig. 1:**
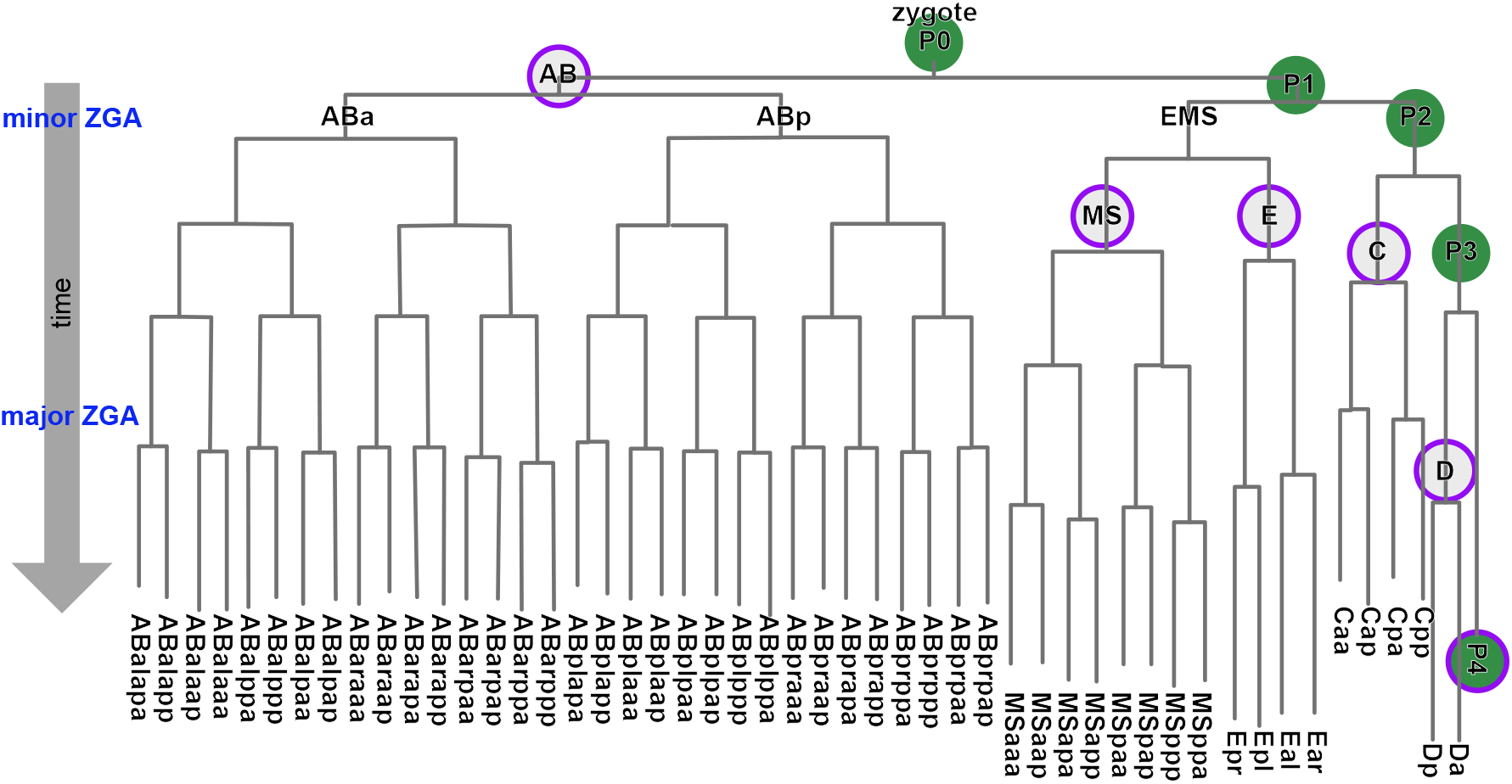
Schematic of the *C. elegans* early embryonic cell lineage^28^, annotated with developmental milestones: minor and major waves of transcription during the zygotic genome activation (ZGA), lineage founder cells from which all other cells of the embryo arise (open purple circles), and germline precursor cells (P-cells) - which are used in CeSCALE as the known anchor cell positions in each embryo.

**Extended Data Fig. 2:**
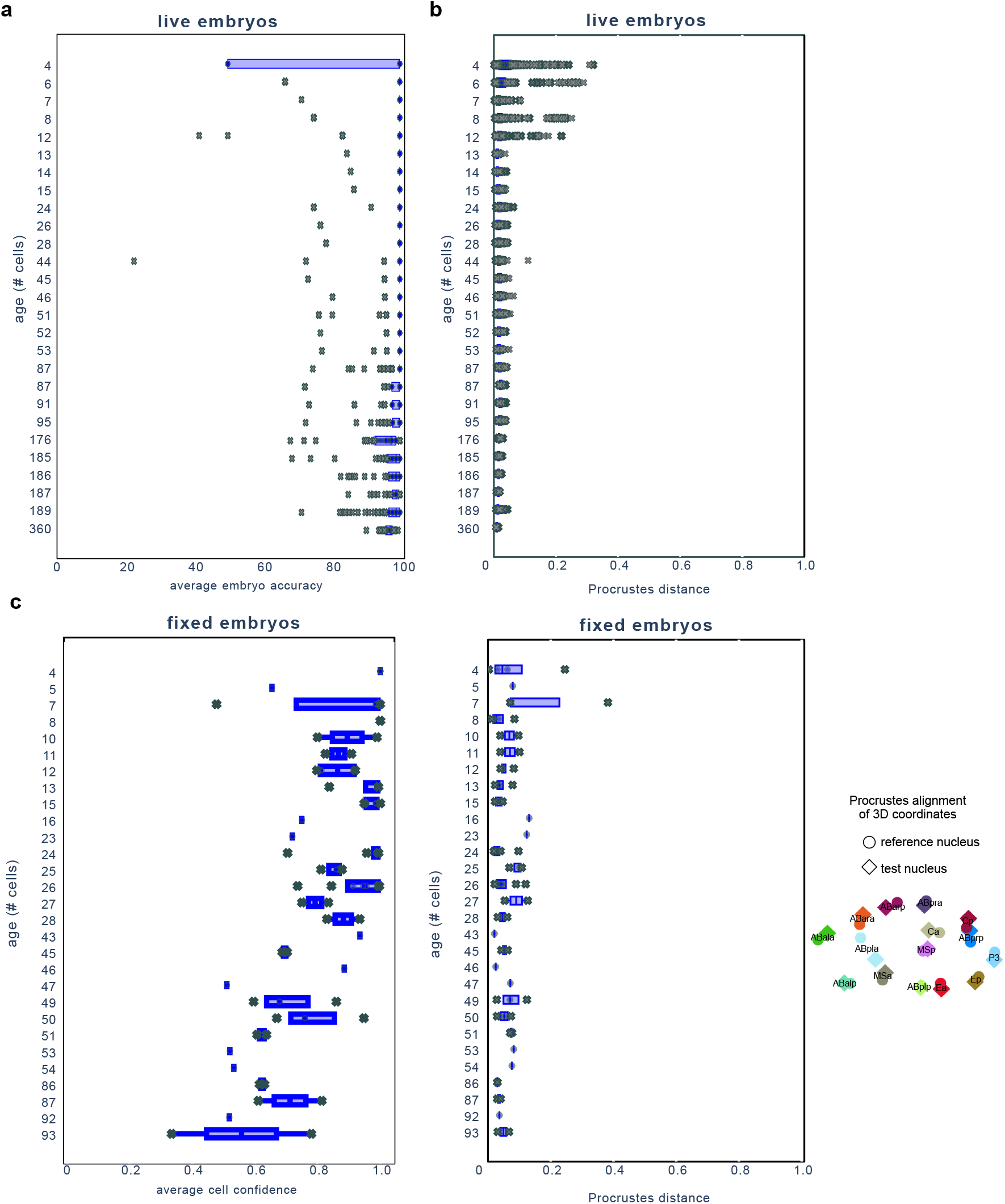
a) Boxplot of accuracy scores for each embryo image stack from live images. The accuracy scores of all nuclei in a given embryo were averaged and all data was plotted by embryo stage. Every data point corresponds to an image. n = 48 embryos, 3923 image stacks. b) (top) Boxplot of Procrustes distances per embryo from live images. n = 48 embryos, 3923 image stacks. (bottom) Boxplot of Procrustes distances per embryo from fixed images. n = 76 embryos, 76 image stacks. Every data point corresponds to an image. An example of superimposed reference and test nuclei positions is shown for a 15-cell embryo. c) Boxplot of confidence values per cell from fixed images. n = 76 embryos, 76 image stacks. Every data point corresponds to an image and the average cell-confidence scores for that image is depicted. All boxplots span Q1–Q3, corresponding to 25th and 75th percentile, respectively, with a line at the median. Any observation below Q1 or above Q3 is plotted as an outlier (× in grey). Whiskers coincide with the edges of Q1 and Q3. See Supplementary Data 4 for individual (average) cell confidence values of ground truth reference cells from live imaging data (representing maximum expected confidence), and Supplementary Data 5 for individual (average) cell confidence values of fixed imaging data.

**Extended Data Fig. 3:**
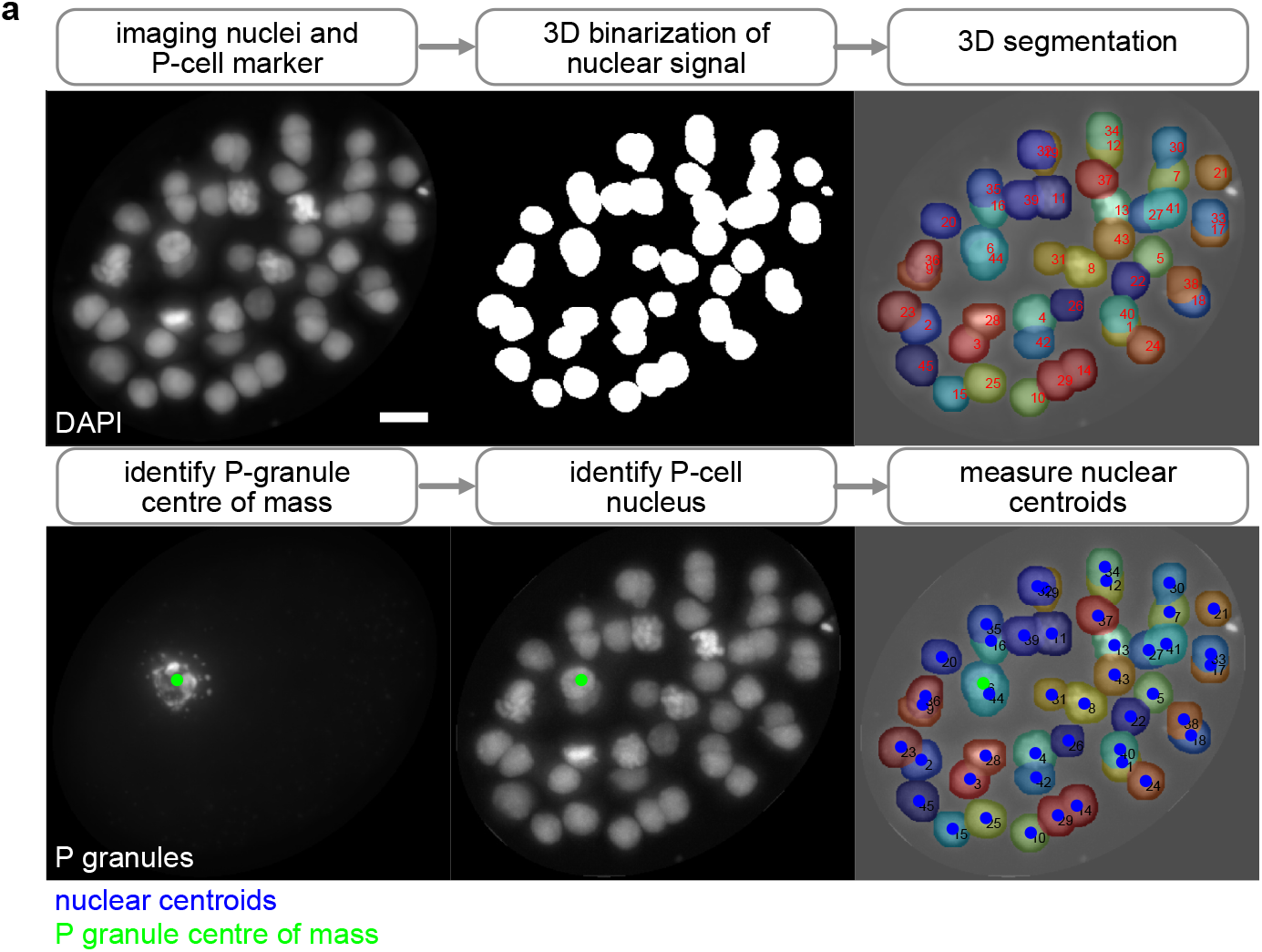
a) Image segmentation method and P-cell identification in fixed samples. 3D image stacks of embryos using DAPI as a nuclear marker, and immunostaining for P-granules which mark the P-cell. Nuclei images are binarized and segmented, the P-granules’ centre of mass is calculated and used to identify the closest nucleus as the P-cell nucleus. Maximum intensity z-projections are displayed. Scale bar, 5 µm.

**Extended Data Fig. 4:**
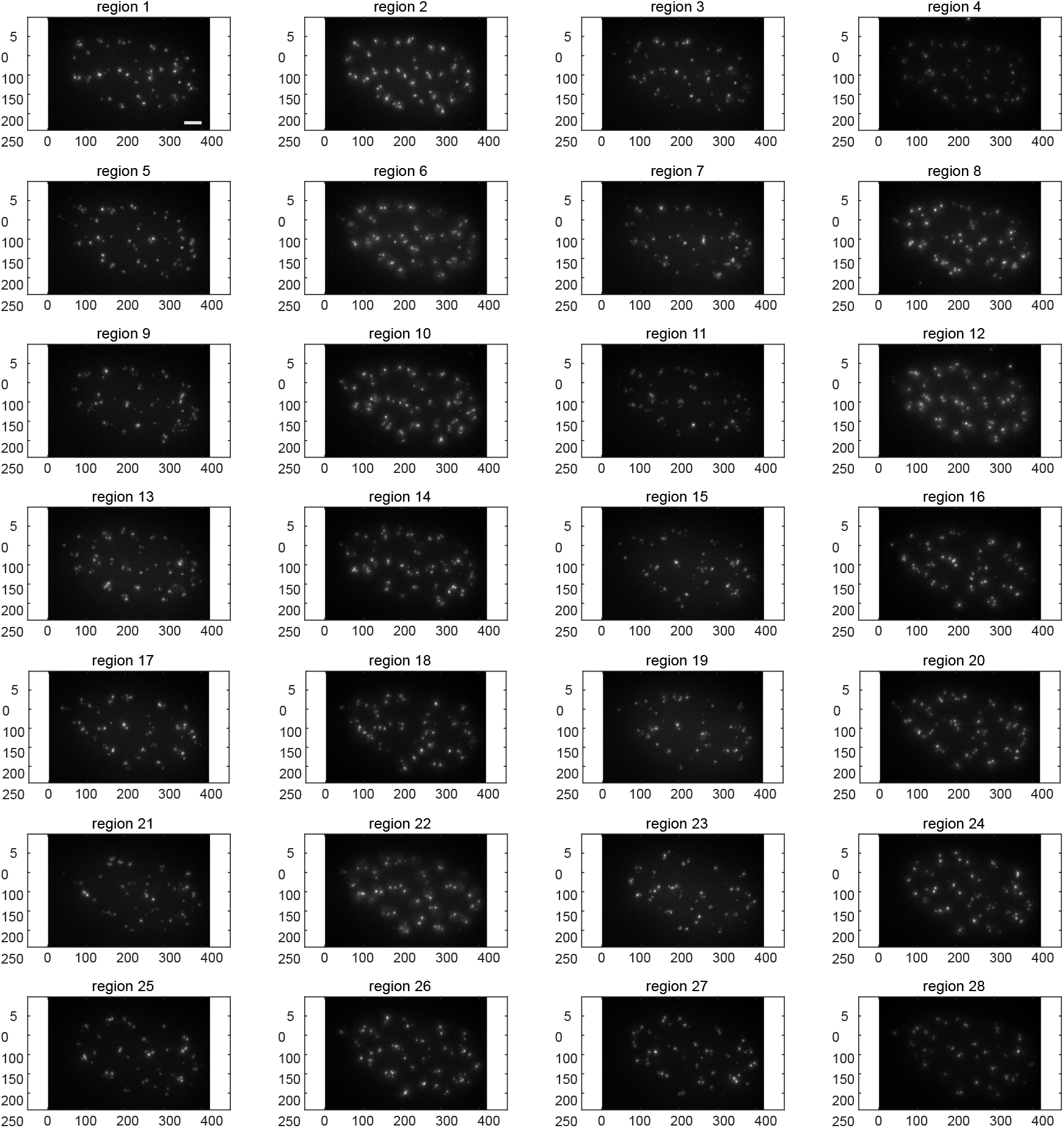
Chromosome tracing secondary hybridizations for chrI and chrII, each dot corresponds to a specific region. Maximum intensity z-projections are displayed.

**Extended Data Fig. 5:**
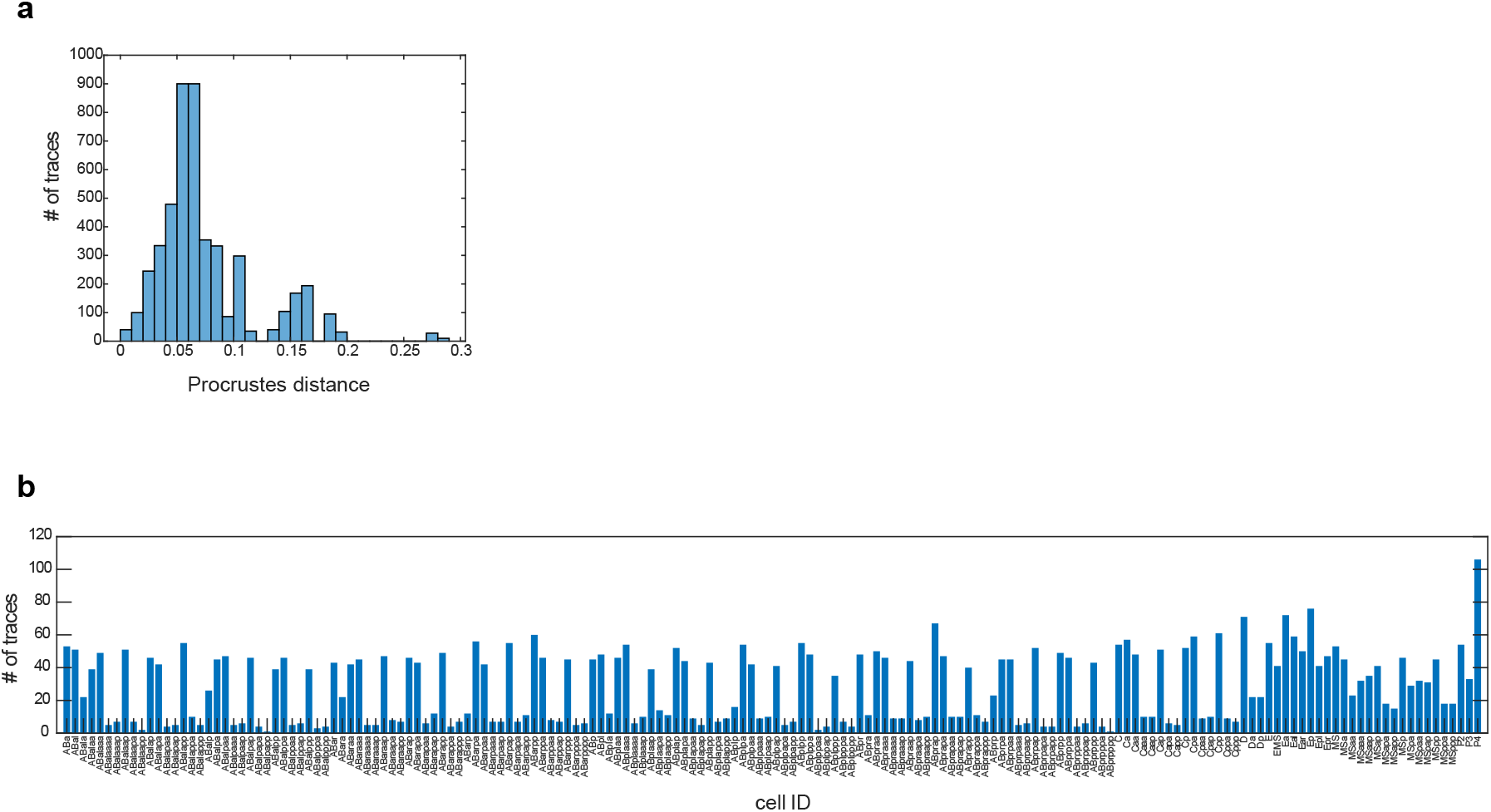
a) Histogram of Procrustes distances per chromosome trace. b) Bar plot of number of chromosome traces detected per cell ID.

**Extended Data Fig. 6:**
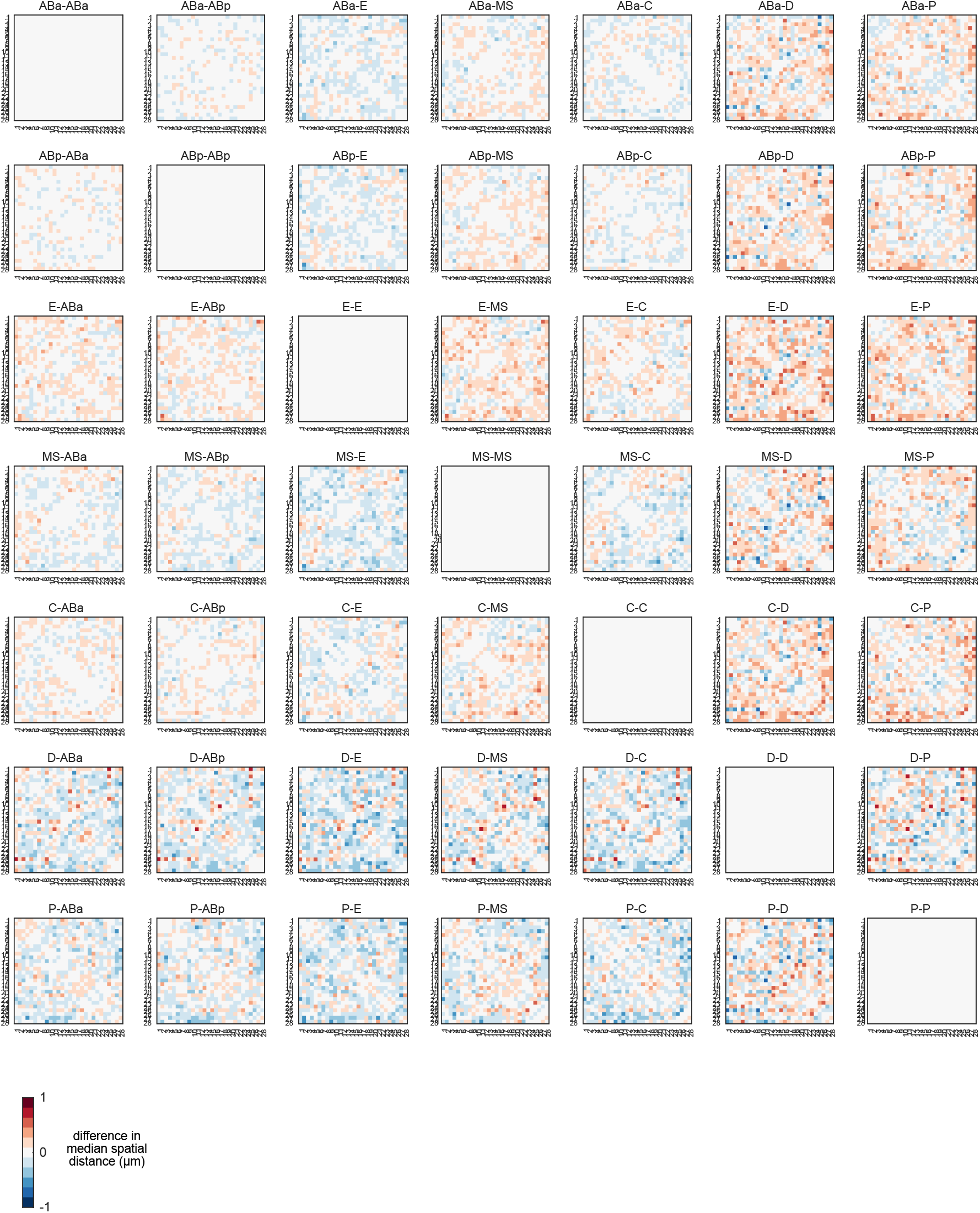
Lineage to lineage difference maps of median chrI and chrII conformation showing global differences in structure.

**Extended Data Fig. 7:**
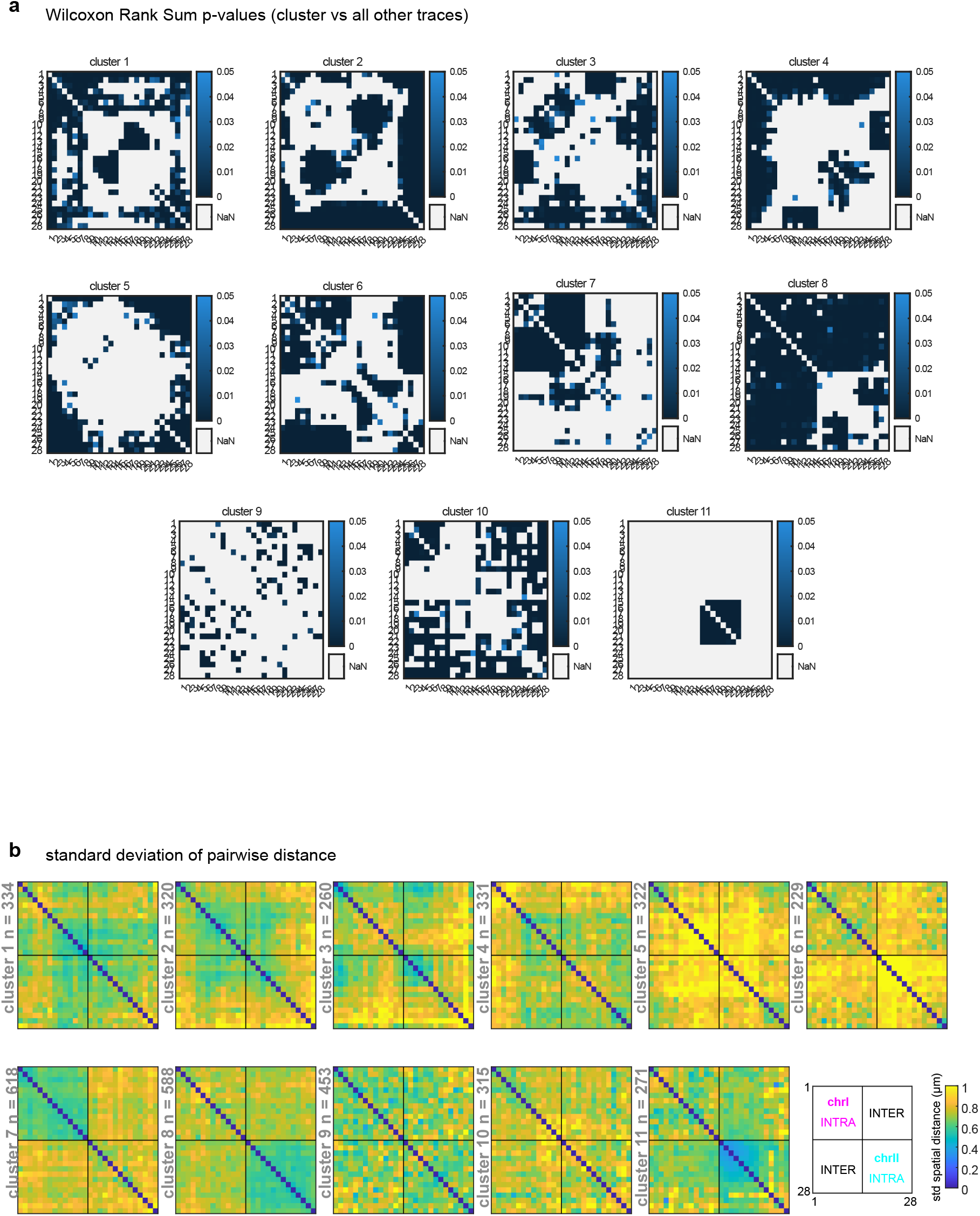
a) P-values of marker regions for each cluster compared to all other chromosome traces, calculated using Wilcoxon Rank Sum Test. b) Standard deviation of spatial distance matrices for each chromosome cluster.

**Extended Data Fig. 8:**
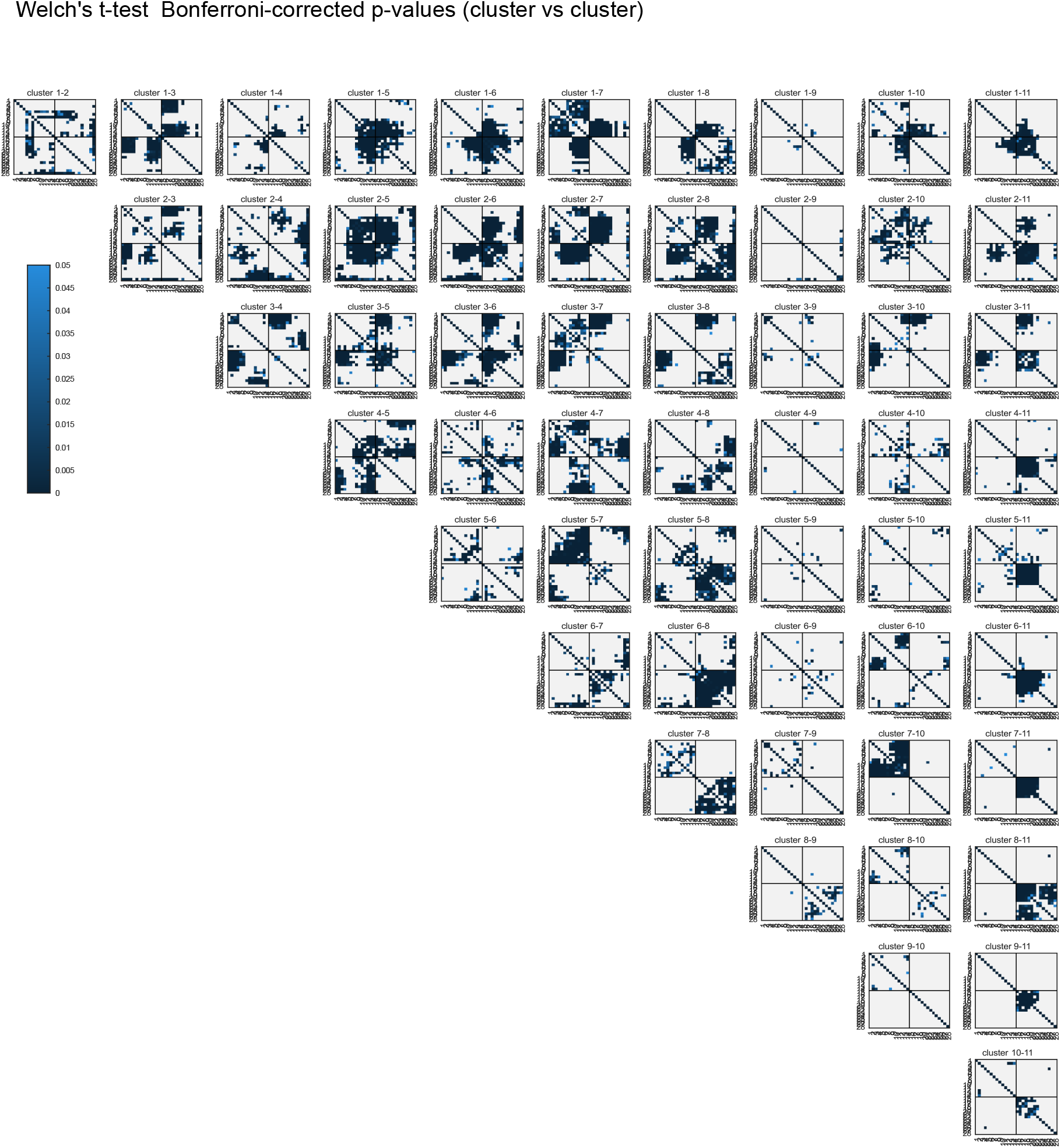
P-values of significantly different pairwise distances between clusters, calculated by Welch’s t-tests, Bonferroni-corrected. **a** all descendents

**Extended Data Fig. 9:**
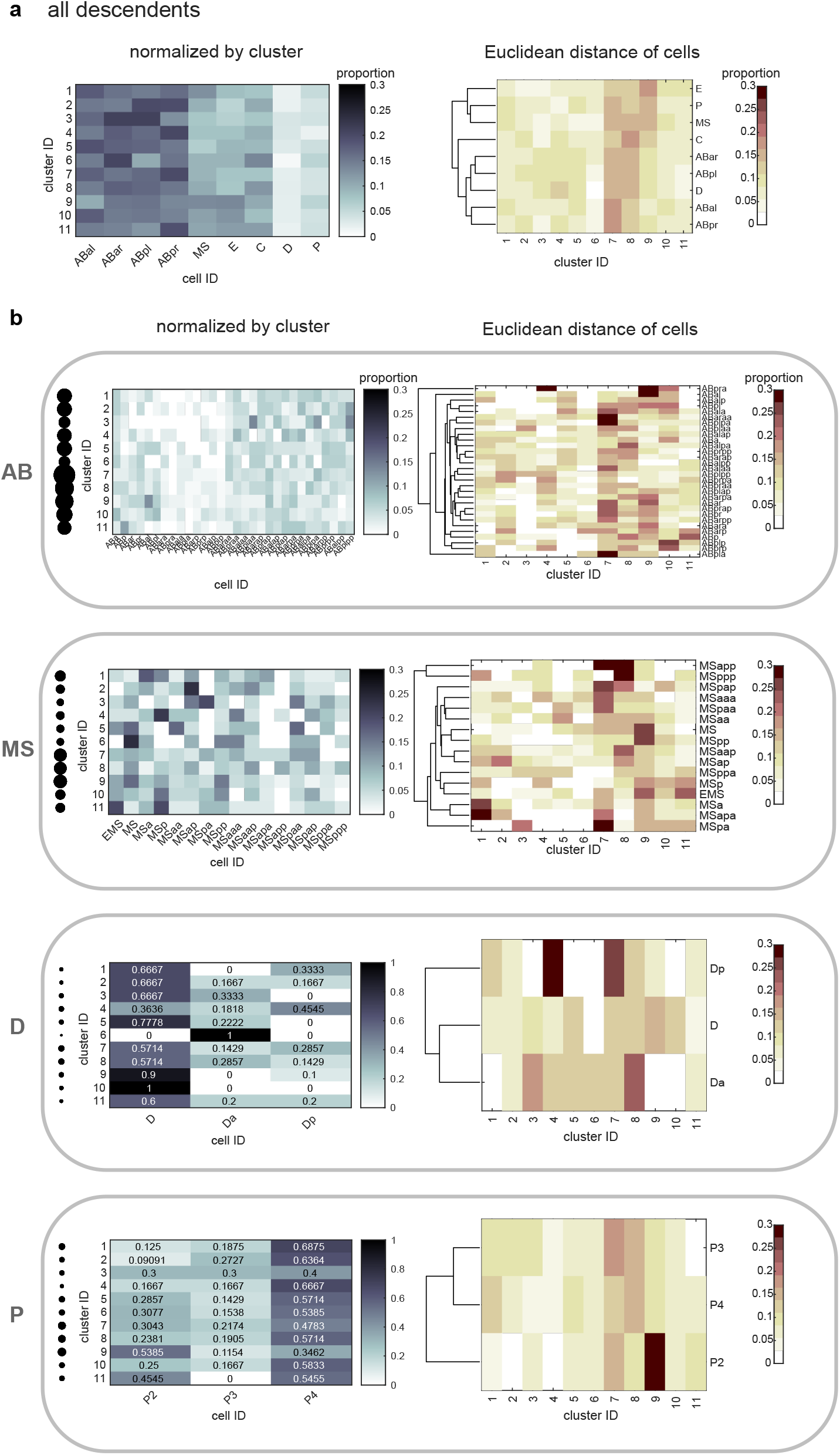
a) Cluster proportions in all cell descendants of each lineage. To determine in which cells each cluster is enriched, each matrix row is normalized to the total amount of traces per cluster (left). Cell proportions in all cell descendants of each lineage. To determine the proportion of clusters within a cell, each matrix column is normalized to the total number of traces per cell ID. The columns of the matrix are ordered using hierarchical clustering by Euclidean distance (right). b) Matrices as in a), parsed by lineage and cell ID in AB, MS, P, and D-cells.

**Extended Data Fig. 10:**
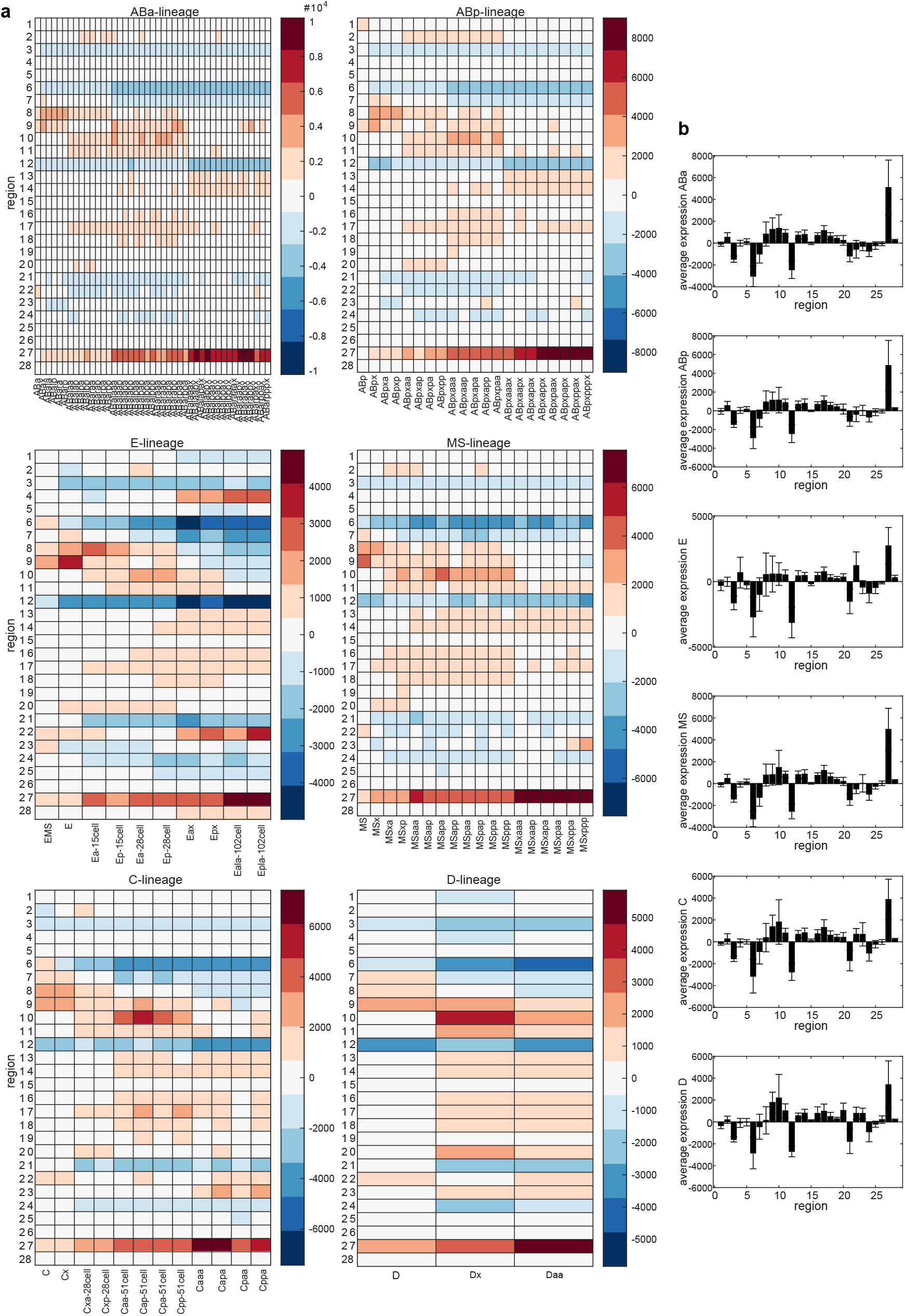
a) Transcript expression heatmaps in chromosome tracing bins per cell. b) Average transcript expression barcharts in chromosome tracing bins per lineage. Error bars represent standard deviation of signal. Units = transcript counts.

**Extended Data Fig. 11:**
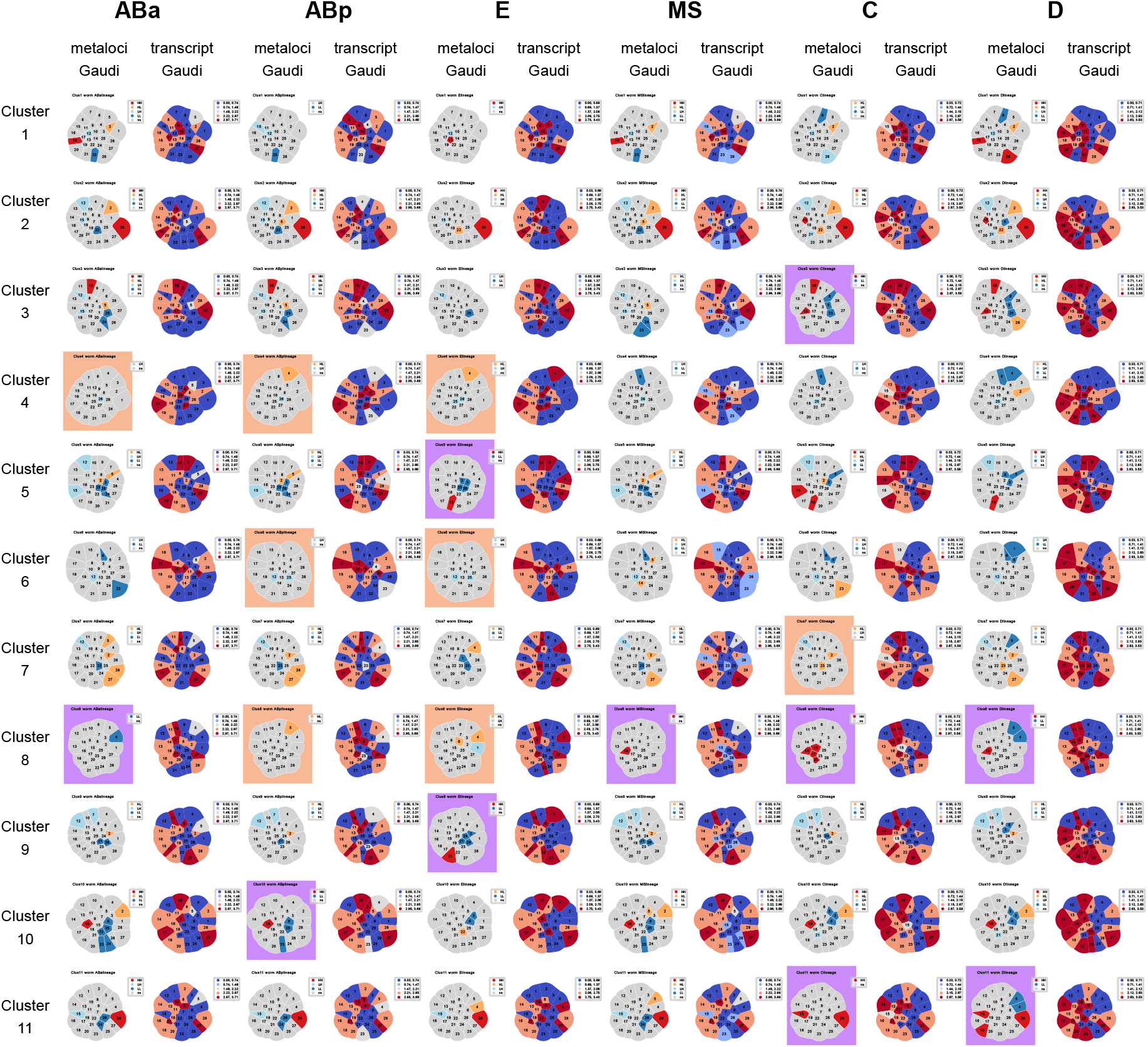
Transcript and Metaloci Gaudi plots for all chromosome clusters in all lineages overall (as in Fig. 5a,c). Structures that contain only HH/LL are highlighted in purple boxes, structures that contain only HL/LH are highlighted in orange boxes.

**Extended Data Fig. 12:**
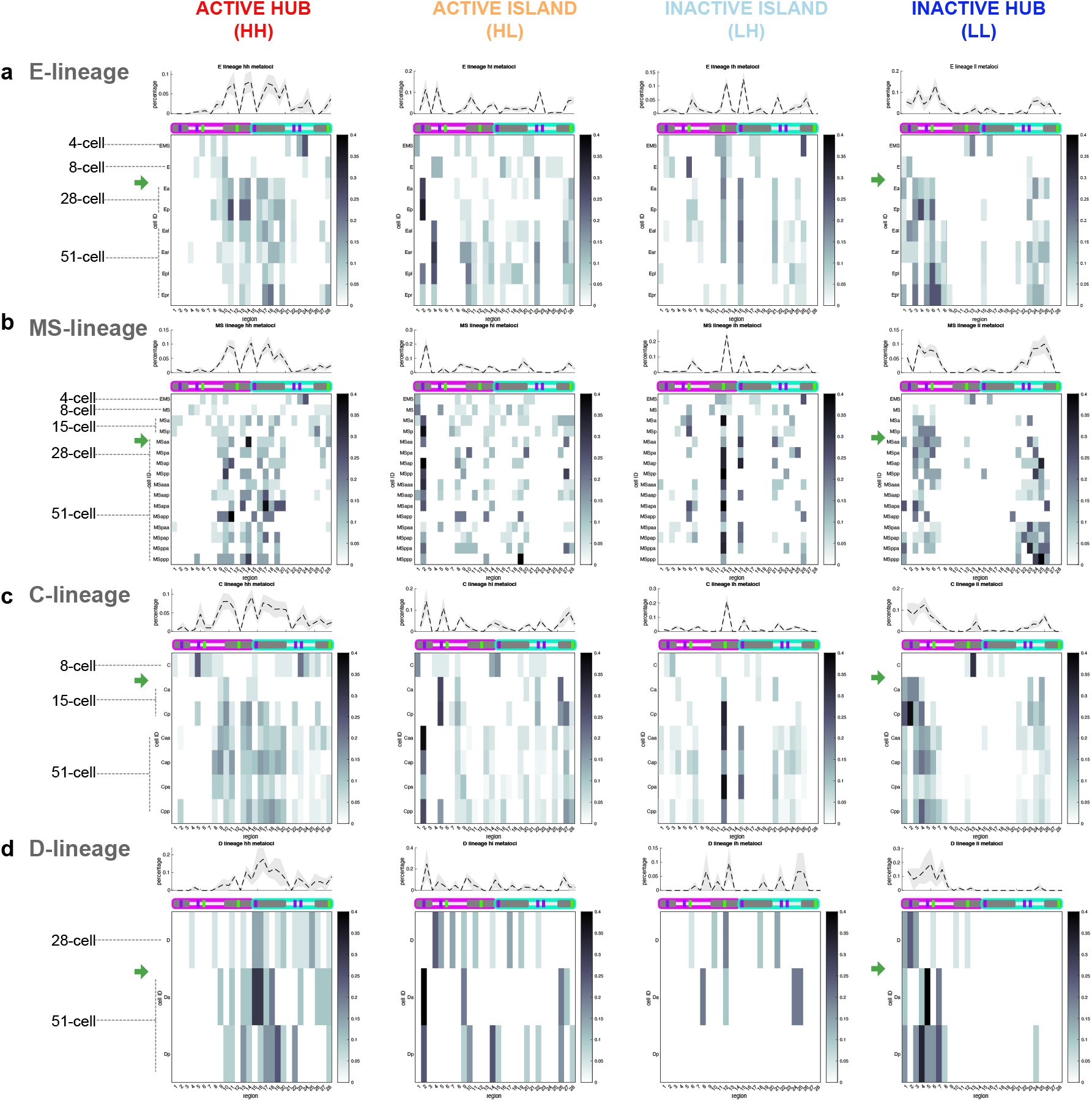
Heatmaps of metaloci proportions in chromosome tracing bins as in Fig. 7, in E, MS, D, and C cells over time.

